# Lifespan Trajectories of Asymmetry in White Matter Tracts

**DOI:** 10.1101/2025.09.29.678806

**Authors:** Sam Bogdanov, Praitayini Kanakaraj, Michael E. Kim, Jessica Samir, Chenyu Gao, Karthik Ramadass, Gaurav Rudravaram, Nancy R. Newlin, Derek Archer, Timothy J. Hohman, Angela L. Jefferson, Victoria L. Morgan, Alexandra Roche, Dario J. Englot, Susan M. Resnick, Lori L. Beason Held, Laurie Cutting, Laura A. Barquero, Micah A. D’Archangel, Tin Q. Nguyen, Kathryn L. Humphreys, Yanbin Niu, Sophia Vinci-Booher, Carissa J. Cascio, The HABS-HD Study Team, Alzheimer’s Disease Neuroimaging Initiative, The BIOCARD Study Team, Zhiyuan Li, Simon N. Vandekar, Panpan Zhang, John C. Gore, Stephanie J. Forkel, Bennett A. Landman, Kurt G. Schilling

**Author notes:** Correspondence to: Sam Bogdanov Vanderbilt University School of Medicine 2209 Garland Avenue, Nashville, TN 37240. HABS-HD MPIs and the HABS-HD Investigators. Data used in preparation of this article were obtained from the Alzheimer’s Disease Neuroimaging Initiative (ADNI) database (adni.loni.usc.edu). As such, the investigators within the ADNI contributed to the design and implementation of ADNI and/or provided data but did not participate in analysis or writing of this report. A complete listing of ADNI investigators can be found at: http://adni.loni.usc.edu/wp-content/uploads/how_to_apply/ADNI_Acknowledgement_List.pdf. These authors contributed equally to this work (Sam Bogdanov, Praitayini Kanakaraj).

## Abstract

Asymmetry in white matter is believed to give rise to the brain’s capacity for specialized processing and is involved in the lateralization of various cognitive processes, such as language and visuo-spatial reasoning. Although studies of white matter asymmetry have been previously documented, they have often been constrained by limited age ranges, sample sizes, or the scope of the tracts and structural features examined. While normative lifespan charts for brain structures are emerging, comprehensive charts detailing white matter asymmetries across numerous pathways and diverse structural measures have been notably absent.

This study addresses this gap by leveraging a large-scale dataset of 35,120 typically developing and aging individuals, ranging from 0 to 100 years of age, from 50 primary neuroimaging studies. We generated comprehensive lifespan trajectories for 30 lateralized association and projection white matter tracts, examining 6 distinct microstructural and macrostructural features of these pathways.

Our findings reveal that: (1) asymmetries are widespread across the brain’s white matter and are present in all 30 pathways; (2) for a given pathway, the degree and direction of asymmetry differ between features of tissue microstructure and pathway macrostructure; (3) asymmetries vary across and within pathway types (association and projection tracts); and (4) these asymmetries are not static, following unique trajectories across the lifespan, with distinct changes during development, and a general trend of becoming more asymmetric with increasing age (particularly in later adulthood) across pathways.

This study represents the most extensive characterization of white matter asymmetry across the lifespan to date, charting how lateralization patterns emerge, mature, and change throughout life. It provides a foundational resource for understanding the principles of white matter organization from early to late life, its relation to functional specialization and inter-individual variability, and offers a key reference for interpreting deviations during healthy development and aging as well as those associated with clinical populations.

## Introduction

Brain asymmetry is an organizing principle of the nervous system that enables functional specialization (Ocklenburg et al., 2024). Nearly two centuries ago, the pioneering clinical observations of Marc Dax (1836) (Dax, 1865; Manning & Thomas-Anterion, 2011) and later Paul Broca (1860s) (Amunts et al., 1999; Broca, 1861) first linked left-hemisphere lesions to specific language deficits, establishing that complex cognitive functions are frequently lateralized to one hemisphere. This hemispheric division of labor is supported by an underlying structural architecture that enhances neural efficiency, allowing for parallel processing of different computations while minimizing cross-hemispheric conduction delays (Brincat & Miller, 2025). Asymmetries in functions as diverse as language (Bishop, 2013), visuospatial attention (Thiebaut de Schotten, Dell’Acqua, et al., 2011), and motor control (Callaert et al., 2011) are thought to depend on this lateralized brain circuitry.

The structural substrate of this functional specialization is the brain’s white matter connectome (Biswal et al., 2010; Sporns, Tononi, & Kotter, 2005). Modern neuroimaging - specifically, diffusion-weighted MRI (dMRI) in combination with fiber tractography (Jeurissen, Descoteaux, Mori, & Leemans, 2019) - enables a noninvasive “virtual dissection” to segment and study the macrostructure of these pathways, as well as a “virtual histology” to probe their tissue microstructure (Alexander, Dyrby, Nilsson, & Zhang, 2019). Macrostructural measures describe morphometric features of pathways size and geometry, including volume, lengths, and areas (Yeh, 2020). In contrast, microstructural measures are sensitive to cellular-level features including axonal coherence, packing density, and myelination (Beaulieu, 2002). Characterizing white matter asymmetry at both macro-and microstructure scales is essential, as they capture distinct biological properties that can follow different trajectories across development and aging.

Many diffusion MRI studies have confirmed robust hemispheric asymmetries in specific white matter tracts, mirroring classic left–right differences in brain function (Dulyan, Bortolami, & Forkel, 2025). In particular, the arcuate fasciculus – a key frontotemporal language pathway – is typically larger or more strongly developed in the left hemisphere, consistent with left-hemisphere dominance for language (Andrulyte et al., 2024). This leftward asymmetry of the arcuate is functionally significant: individuals with a more pronounced leftward arcuate show stronger left-hemisphere activation during speech processing tasks (Ocklenburg, Hugdahl, & Westerhausen, 2013; Propper et al., 2010), and higher performance in verbal tasks (Catani et al., 2007). Conversely, frontoparietal pathways associated with attention tend to be right-lateralized – for example, the inferior branch of the superior longitudinal fasciculus (SLF-III) is consistently larger on the right (Amemiya, Naito, & Takemura, 2021; Howells et al., 2018; Thiebaut de Schotten, Dell’Acqua, et al., 2011), aligning with the right hemisphere’s specialization in spatial attention. Notably, these tract-level asymmetries emerge by early childhood and are thought to scaffold normal cognitive development (Ghasoub et al., 2025; Marcelle, Illapani, Gaillard, & Newport, 2025) (e.g. supporting mature language networks). Further, atypical patterns of white matter laterality have been linked to clinical conditions, with altered or atypical (or even reversed) patterns of white matter asymmetry reported in schizophrenia (Gomez-Gastiasoro et al., 2019; Ho et al., 2017), autism spectrum disorder (Carper, Treiber, DeJesus, & Müller, 2016), dyslexia (Banfi et al., 2019; Rimrodt, Peterson, Denckla, Kaufmann, & Cutting, 2010), and Parkinson’s Disease (Chen, Deng, Zuo, & Zhong, 2022), among others (Saltoun, Yeo, Paul, Diedrichsen, & Bzdok, 2025).

Despite this growing body of work (Carper et al., 2016; Demnitz et al., 2021; Hau et al., 2017; Honnedevasthana Arun, Connelly, Smith, & Calamante, 2021; V. Kumpulainen et al., 2023; Mundorf, Peterburs, & Ocklenburg, 2021; Roe et al., 2023; M. Shu et al., 2021; N. Shu, Liu, Duan, & Li, 2015; Takao, Hayashi, & Ohtomo, 2013; Y. Zhu et al., 2023), current knowledge of white-matter asymmetry rests largely on static snapshots from studies with modest sample sizes, narrow age ranges, and limited pathway and feature coverage. This has led to a fragmented literature in which the reported laterality for a given pathway often diverges, due to variations in sample size, age range/cohorts, and the specific structural feature(s) investigated. Moreover, most reports simply focus on testing whether the group mean differs from zero, rather than describing the age-specific distribution and how it changes across development and aging. This leaves unanswered questions about when asymmetry emerges, whether it strengthens or weakens at different life stages, and how widely it varies among healthy individuals at a given age. In short, the field lacks a lifespan-wide normative reference that quantifies the distribution of tract-level asymmetry across development and aging.

Here, we assemble 35,120 scans from 50 population-based cohorts spanning 0-100 years and construct age-varying brain asymmetry charts for white-matter pathways. Rather than centering on a binary test of whether asymmetry exists, we use an estimation framework to quantify the *magnitude*, *direction*, and *age-specific distribution* of asymmetry. Specifically, we: (1) establish normative lifespan trajectories for all investigated features and pathways; (2) identify which tracts and features exhibit asymmetry during key periods of development, adulthood, and aging; (3) quantify the population-level distribution of right–left asymmetries within these periods; (4) determine whether - and at what ages - the direction of asymmetry reverses; and (5) assess how the magnitude of asymmetry changes dynamically (i.e., increases or decreases) across infancy, development, and later life. A priori, we anticipate that normative modelling would reveal structured, tract-and feature-specific distributions of laterality that change across the lifespan, including shifts in both the median and the spread of asymmetry across development and aging. Together, these charts provide a population reference to contextualize individuals and cohorts, enabling future developmental and clinical studies to interpret asymmetry in an age-appropriate framework.

## Materials and Methods

We analyzed 35,120 diffusion MRI scans from 50 population-based cohorts spanning 0-100 years (**Figure 1A**). For each participant, we derived tract-level measures for 30 bilateral white-matter pathways, capturing both microstructure (diffusion tensor imaging indices reflecting tissue organization, axonal density, and myelination) and macrostructure (tract volume and length) (**Figure 1B**). We computed a laterality index for every tract-feature pair (**Figure 1C**) and modelled age-varying centile curves to estimate the population distribution (i.e. typically showing 2.5th-25th-50th-75th-97.5th centiles) at each age. Our analyses summarized patterns across the developmental and aging windows, quantify inter-individual variability, and identify tracts and features with asymmetry or direction reversals.

**Figure 1.**
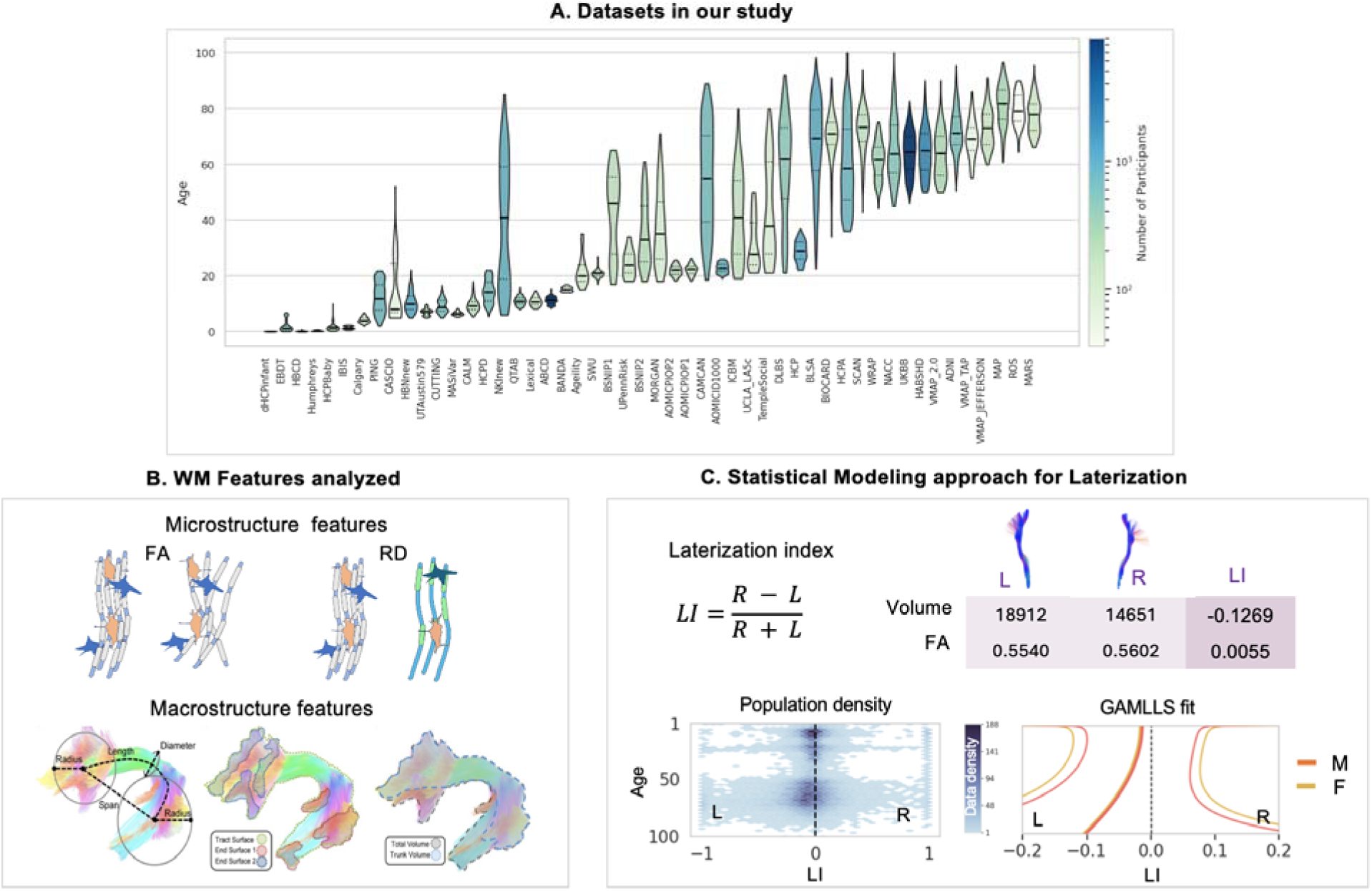
Overview of the study datasets, white matter features, and analytical framework. (**A**) Age distributions for each of the 50 contributing datasets (violin plots), illustrating broad coverage from 0–100 years. Color encodes the number of participants per dataset (log scale). (**B**) Features extracted for each of the 30 bilateral pathways. Microstructural indices (e.g., Fractional Anisotropy, and Mean, Axial and Radial diffusivities; FA, MD, AD, RD) summarize tissue organization and axonal/myelin density; macrostructural indices (e.g., tract volume, length, surface area) capture pathway size and geometry. (**C**) Analysis pipeline. For each participant, white matter pathways were segmented, and features were extracted. A Lateralization Index (LI) was calculated for each tract-feature pair. These LIs were used as input for a normative modeling framework (GAMLSS) to generate age-specific population centile curves.

We note that we emphasize and describe population *centiles* - which describe the expected range of individual values at a given age - rather than *confidence intervals*, which only quantify the precision of the median estimate; we therefore do not perform null-hypothesis significance tests, which are trivially positive at very large N. Study cohorts, image preprocessing, white matter pathway segmentation, statistical analysis and normative modeling are described below.

### Participants and Study Cohorts

The final dataset comprised 35,120 cross-sectional diffusion MRI scans aggregated from 50 independent, population-based cohorts (**Figure 1A**; see **Supplementary Table 1** for a detailed description of each cohort). To ensure a cross-sectional analysis, only the earliest available scan was included for any participant with longitudinal data, so that each individual is represented only once. The cohorts consist of typically developing and aging participants with broad geographic representation. Studies were included which contained broad patient populations; for these studies, only neurotypical participants without a pathologic diagnosis (as defined by each cohort) were included in the analyses performed in this study. Acquisition parameters varied across studies (see **Supplementary Table 2** for detailed description of voxel sizes and b-values), necessitating downstream harmonization - including both standardized preprocessing (see *Diffusion MRI Processing Pipeline*), unform tract segmentation (see *Tract Segmentation*), and statistically within the lifespan modelling (see *Lifespan Normative Modeling Framework*). Demographic information was harmonized across studies where available, including sex (53.7% female) and handedness (of 14,220 participants with handedness data, 9.3% were left-handed). In studies with handedness available, self-report or a questionnaire (typically the Edinburgh Handedness Inventory), were used to determine handedness (handedness experiments are given as supplementary material). Cognitive and behavioral measures were not consistently available across the consortium and were therefore not included in the present analysis. All contributing studies received ethical approval from their local institutional review boards.

### Diffusion MRI Processing Pipeline

All diffusion MRI data were processed with a single, standardized workflow using the PreQual pipeline (v 1.0.8) applied identically across cohorts to promote reproducibility (Cai et al., 2021). This pipeline corrected for susceptibility-induced EPI distortions, head motion, and eddy current artifacts. To ensure consistency and reproducibility across the 50 cohorts, which varied in scanner and acquisition parameters, the same pipeline version was applied to all datasets, with specific flags adapted for each study’s raw data structure. Following this preprocessing, diffusion tensor imaging (DTI) models were fitted to generate voxel-wise maps of fractional anisotropy (FA), mean diffusivity (MD), axial diffusivity (AD), and radial diffusivity (RD). Quality control was conducted at two stages (Michael E Kim, Gao, et al., 2024; Michael E Kim, Ramadass, et al., 2024). First, automated logs and visual reports from PreQual were reviewed to identify corruption, gross motion, or failed corrections. Second, the derived scalar maps were visually inspected to verify anatomical plausibility and artifact mitigation. Scans failing QC at either stage were excluded prior to downstream analyses.

### White Matter Pathway Segmentation and Feature Extraction

#### Tract Segmentation

The preprocessed dMRI data for each participant was upsampled to 1 mm isotropic resolution and input into TractSeg (v2.8),(Wasserthal, Neher, & Maier-Hein, 2018) a convolutional neural network (CNN) framework for white matter bundle segmentation.(Wasserthal et al., 2018) TractSeg operates directly on local fiber orientation images, segmenting tracts without requiring whole-brain tractography or inter-subject registration. We employed the TractSeg pipeline with default parameters, which uses the MRtrix3 implementation of Constrained Spherical Deconvolution (CSD) to extract fiber orientation peaks from the DWI data for downstream bundle segmentation, bundle endpoints segmentation, tract-orientation mapping, and subsequent tractography. The pre-trained model automatically segmented 72 white matter bundles per participant. From these, we selected 60 lateral pathways (i.e., 30 bilateral tract pairs), excluding midline commissural structures, for asymmetry analyses. The set spans association and projection systems (with limbic, thalamic, and striatal subdivisions noted in **Table 1**).

**Table 1.**
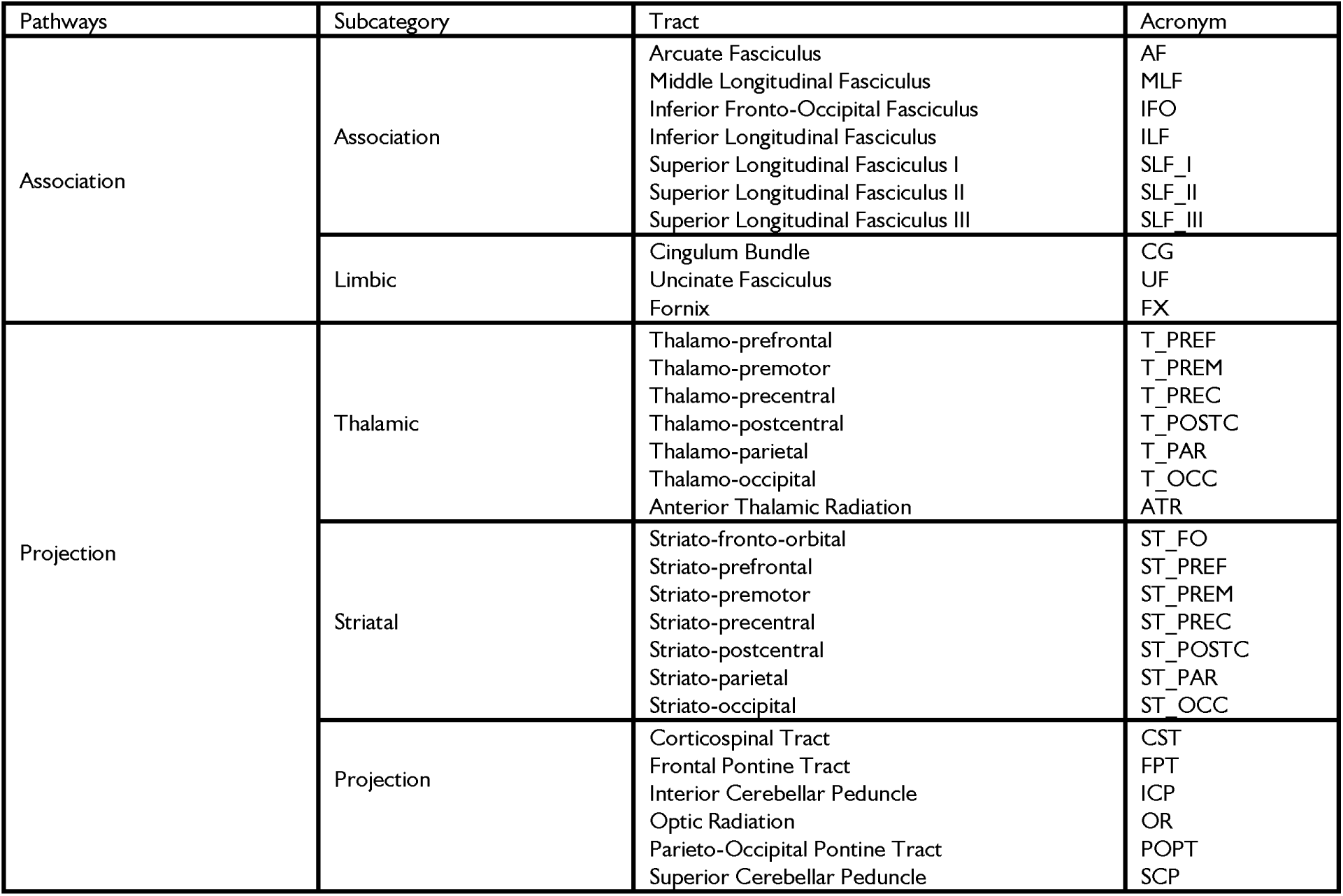
List of 30 Bilateral White Matter Tracts Included in Asymmetry Analysis. As defined by the preset anatomical labels and segmentation outputs of the automated tractography tool TractSeg.

#### Microstructural and Macrostructural Feature Definition

For each of the 30 segmented tract pairs, we computed two classes of features using the scilpy library (Sherbrooke Connectivity Imaging Lab’s open-source toolkit, version 1.5.0)(scilus/scilpy) (Figure **1B**).

Microstructural Features: We calculated four standard DTI indices that reflect cellular-level tissue properties (Beaulieu, 2002; Wheeler-Kingshott & Cercignani, 2009). To derive a single value for each tract, we computed a weighted average of the metric across all voxels within the tract’s mask, with the contribution of each voxel weighted by the number of streamlines passing through it.

- Fractional Anisotropy (FA): an index of directional coherence of diffusion, influenced by axonal organization/packing and myelin sheaths.
- Mean Diffusivity (MD): the average magnitude of water diffusion, sensitive to overall water mobility within tissue, influenced by axonal/myelin densities.
- Axial Diffusivity (AD): magnitude of water diffusion parallel to the principal fiber direction sensitive to the intra-axonal space and changes in axonal caliber and architecture.
- Radial Diffusivity (RD): magnitude of water diffusion perpendicular to the principal fiber direction. As myelin sheaths are a primary barrier to perpendicular diffusion, increased RD is often interpreted as reflecting demyelination or reduced axonal packing.

Macrostructural Features: We calculated three features describing the large-scale geometry and morphology of each tract (Schilling et al., 2023; Yeh, 2020):

- Tract Volume: The total volume in mm³ occupied by the pathway.
- Mean Streamline Length: The average length in mm of the streamlines constituting the tract, reflecting the trajectory extent.
- Tract Surface Area: The surface area in mm² of the geometric shape enclosing the tract, indexing bundle envelope size/complexity.

### Statistical Analysis and Normative Modeling

#### Calculation of the Lateralization Index (LI)

For each micro-and macrostructural feature, we calculated a *Lateralization Index* (LI) to quantify the degree of asymmetry between the left (L) and right (R) hemispheres for each tract (**Figure 1C**). We used the formula 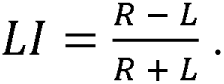 This index is bounded between-1 and 1. An LI of 0 indicates perfect symmetry. For this study, positive LI values indicate a rightward asymmetry (i.e., a higher value in the right-hemisphere tract), while negative LI values indicate a leftward asymmetry (a higher value in the left-hemisphere tract) (Savic & Lindström, 2008). This definition (and data presentation) of the lateralization index is consistent with the expert consensus provided in the *Laterality indices consensus initiative* (Vingerhoets et al., 2023).

#### Lifespan Normative Modeling Framework

We created lifespan charts of white matter asymmetry using a normative modeling approach. In the context of this large-scale dataset (N > 35,000), traditional null-hypothesis significance testing is not informative, as even biologically negligible asymmetries would yield a statistically significant result. Our goal was therefore not to ask the binary question of *whether* asymmetry exists, but rather to adopt an estimation framework to describe the full, age-specific distribution of the LI. This approach, analogous to the creation of pediatric growth charts (Grummer-Strawn, Reinold, Krebs, Centers for Disease, & Prevention, 2010; Santoni et al., 2020), allows for the characterization of the typical range of inter-individual variation at any given age (Bethlehem, Seidlitz, et al., 2022).

To model these complex, non-linear lifespan trajectories, we employed Generalized Additive Models for Location, Scale, and Shape (GAMLSS), implemented in R using the gamlss package (Bethlehem, Seidlitz, et al., 2022; Rigby & Stasinopoulos, 2005). GAMLSS is highly suited for lifespan data as it flexibly models how the entire distribution of the LI - including its median (location, µ), variability (scale, σ), and shape - changes as a non-linear function of age.

Diffusion MRI measures can vary systematically across scanners and acquisition protocols in multi-cohort datasets (Grech-Sollars et al., 2015; Kurt G. Schilling et al., 2021). To mitigate these effects, we modeled study cohort as a dataset-specific effect on both the location and scale of the LI distribution as in (BethlehemSeidlitz, et al., 2022), allowing cohort-level offsets and dispersion differences while estimating a shared age-varying trajectory. In addition, the LI is a within-subject normalized ratio (R−L)/(R+L), which reduces sensitivity to global scaling differences that affect left and right measurements similarly within an individual.

For each of the 30 tracts and 7 features, a separate GAMLSS model was fit. We modeled the LI as a response variable following a normal distribution (Savic & Lindström, 2008), where both the mean (µ) and the standard deviation (σ) were modeled as functions of age, sex, and study cohort:

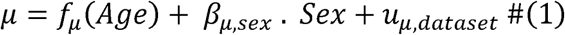

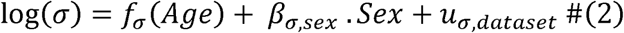

Here, fμ(⋅) and fσ(⋅) are smooth functions of age, sex is a fixed effect, and dataset random effects (u_μ_,u_σ_) capture residual between-study variation. Allowing σ to vary with age accommodates age-dependent heteroscedasticity.

Age effects were represented with fractional polynomial (FP) smooths (order up to 3; powers drawn from [-2, −1, −0.5, 0, 0.5, 1, 2, 3], with logarithmic terms for repeated powers). For each tract–feature we fit candidate models with different FP choices for μ and σ and selected the specification with the lowest Bayesian Information Criterion (BIC) (Bethlehem, Seidlitz, et al., 2022).

From the parameters of the best-fitting model, we generated smooth centile curves (e.g., 2.5th, 25th, 50th, 75th, and 97.5th percentiles) across a dense age grid, forming the final normative brain asymmetry charts (**Figure 1C**).

#### Characterizing Lifespan Dynamics

To examine white matter asymmetry across the lifespan, we used the normative LI trajectories derived from our GAMLSS models and summarized them using six predefined lifespan epochs: early childhood (2–5 years), childhood (5–12 years), adolescence (12–20 years), early adulthood (20–40 years), midlife (40–60 years), and older adulthood (60–100 years). For analyses requiring discrete “snapshot” summaries, we selected six representative ages corresponding to the median age within each epoch (3, 9, 17, 30, 50, and 80 years). For window-based analyses, we quantified trends within each epoch directly.

1. What are the characteristics of exemplar lifespan asymmetry curves?

To answer this, we visualized the full, sex-specific centile curves (2.5th to 97.5th percentile) for selected white matter tracts and features to illustrate general patterns of development, aging, and inter-individual variability.

2. Which features and pathways exhibit population-level asymmetry at critical lifespan stages?

We summarized tract-and feature-specific asymmetries at six representative ages (3, 9, 17, 30, 50, and 80 years) using heatmaps that display both the population median LI and the proportion of individuals with rightward vs. leftward lateralization.

3. Do asymmetries reverse direction across the lifespan?

We identified shifts in hemispheric dominance by assessing zero-crossings in the median LI trajectories, which indicate if and when the direction of population-level asymmetry changed (e.g., from leftward to rightward, or vice versa).

4. How does the magnitude of asymmetry evolve during different life stages?

Within each of the six lifespan epochs, we quantified the rate of change in asymmetry magnitude by calculating the mean slope of the absolute value of the median LI curve. This allowed us to determine whether the *magnitude of asymmetry* was increasing (strengthening) or decreasing (attenuating) during that life stage.

## Results

We constructed normative lifespan charts of white matter asymmetry by applying our statistical modeling framework to 35,120 individuals spanning 0-100 years. These charts characterize the magnitude, direction, and population-level variability for 6 micro-and macrostructural features across 30 bilateral white matter pathways. This comprehensive analysis revealed several key principles of brain lateralization, which are detailed in the following sections. We begin by illustrating the modeling approach and showing exemplar trajectories that highlight distinct patterns of development and aging.

### What are the characteristics of exemplar lifespan asymmetry curves?

The normative lifespan charts revealed that white matter tracts exhibit highly heterogeneous and dynamic patterns of asymmetry, which varied substantially across different pathways, features, and ages. To orient the reader to these results, **Figure 2** illustrates the full centile-based lifespan trajectory for two features of the arcuate fasciculus (AF) – the FA and total volume. These charts depict the population median (50th percentile) and the spread of the population (from the 2.5th to the 97.5th percentile), with lateralization intuitively shown left-to-right along the x-axis, providing a comprehensive view of both the central tendency and the inter-individual variability in asymmetry at any given age.

**Figure 2.**
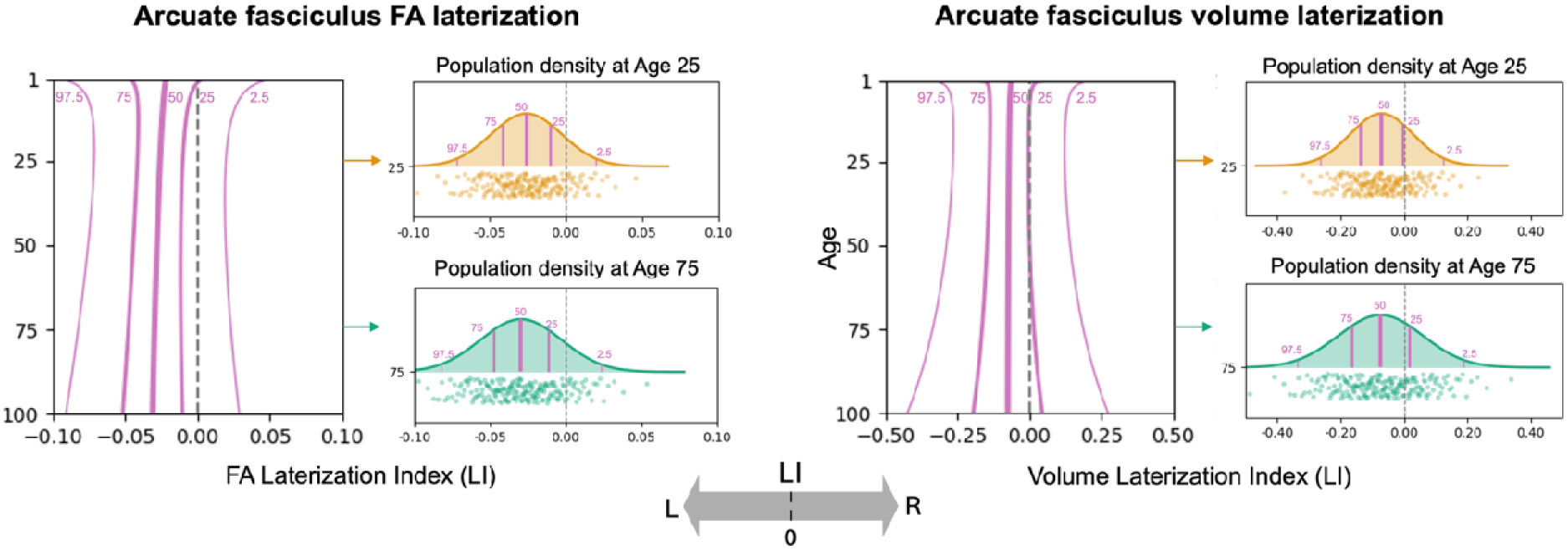
Exemplar asymmetry lifespan curves for the arcuate fasciculus (AF). Lifespan trajectories for fractional anisotropy (FA; a microstructural feature) and tract volume (a macrostructural feature) are shown. Lines represent the 2.5th, 25th, 50th (median), 75th, and 97.5th centile curves, illustrating the full population distribution of the Laterality Index (LI) across the lifespan. Negative values indicate leftward asymmetry, positive values indicate rightward asymmetry.

Building on this, **Figure 3** displays a broader selection of exemplar trajectories for four well-studied pathways (AF, Anterior Thalamic Radiation [ATR], Frontal Pontine Tract [FPT], Corticospinal Tract [CST]) across six key micro-and macrostructural (FA, MD, AD, RD, tract volume, tract length) features. Analysis of asymmetry in these same pathways and features according to handedness was also performed (**Supplementary Figure 1**).

**Figure 3.**
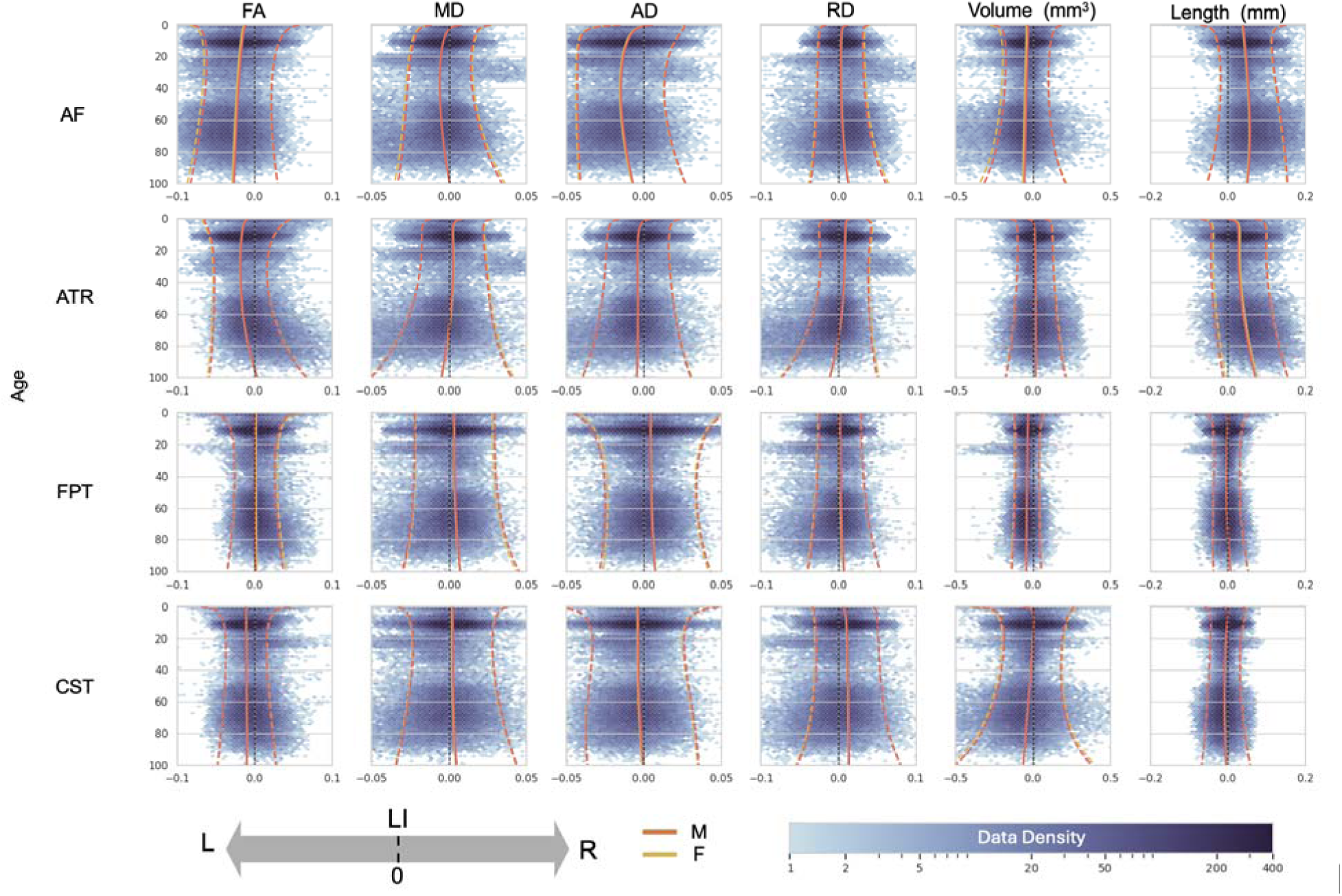
Diverse patterns of lifespan asymmetry across tracts and features. Lifespan trajectories of the LI are shown for four white matter tracts (arcuate fasciculus (AF), anterior thalamic radiation (ATR), frontal pontine tract (FPT), and corticospinal tract (CST)) and six features (FA, MD, AD, RD, volume, and length). Solid lines represent the population median (50th percentile), with the lighter bands showing the inter-quartile range (25th-75th percentiles). Red and yellow lines correspond to males and females, respectively. The plots highlight that the magnitude, direction, and age-related dynamics of asymmetry are highly specific to the tract and feature being measured. Full charts for all tracts and features are provided in **Supplementary Material (Supplementary Fig. S2)**.

These charts highlight several overarching principles: First, while asymmetry is common, its magnitude varies by feature type. Microstructural asymmetries were often subtle, with the bulk of the population having LI values between-0.05 and +0.05. For example, the AF showed a consistent leftward FA asymmetry with a population median LI ranging from-0.02 to-0.03 across age and 25^th^/75^th^ centiles from-0.05 to-0.01. In contrast, macrostructural features exhibited a much wider range of individual variability, often spanning the range from-0.2 to +0.2, with larger median effects (e.g., median LI for ATR length ∼0.05; FPT volume ∼-0.07). Second, patterns were highly dependent on the combination of tract and metric, where pathways can show opposite lateralization directions across features, highlighting the need to study both micro-and macrostructure jointly. For example, the AF shows left-lateralization for FA and volume but right-lateralization for RD and length, or the ATR showing general leftward asymmetries for diffusivities and rightward volume and length asymmetries. Third, these asymmetries were not static but changed dynamically across the lifespan, with many tracts showing age-dependent increases or decreases in lateralization. Fourth, despite these complex dynamics, male and female median trajectories showed general overlap across nearly all tracts and features, suggesting minimal sex differences in the average patterns of asymmetry. Finally, inter-individual variability was not constant, with wider centile bands in early childhood and older age suggesting a greater diversity of asymmetry patterns during these life stages.

All individual lifespan charts for all 30 tracts and 6 features are provided in the **Supplementary Material (Supplementary Fig. 2)**.

### Which features and pathways exhibit population-level asymmetry at critical lifespan stages?

To characterize asymmetry at discrete life stages, we examined the population median LI (**Figure 4**) and the percentage of individuals with rightward lateralization (**Figure 5**) at six key ages: 3, 9, 17, 30, 50, and 80 years (representative of early childhood, childhood, adolescence, early adulthood, midlife, and older adulthood, respectively). Together, the two figures convey both the typical direction/magnitude (median LI) and the population consensus (prevalence) for each tract–feature–age combination.

**Figure 4.**
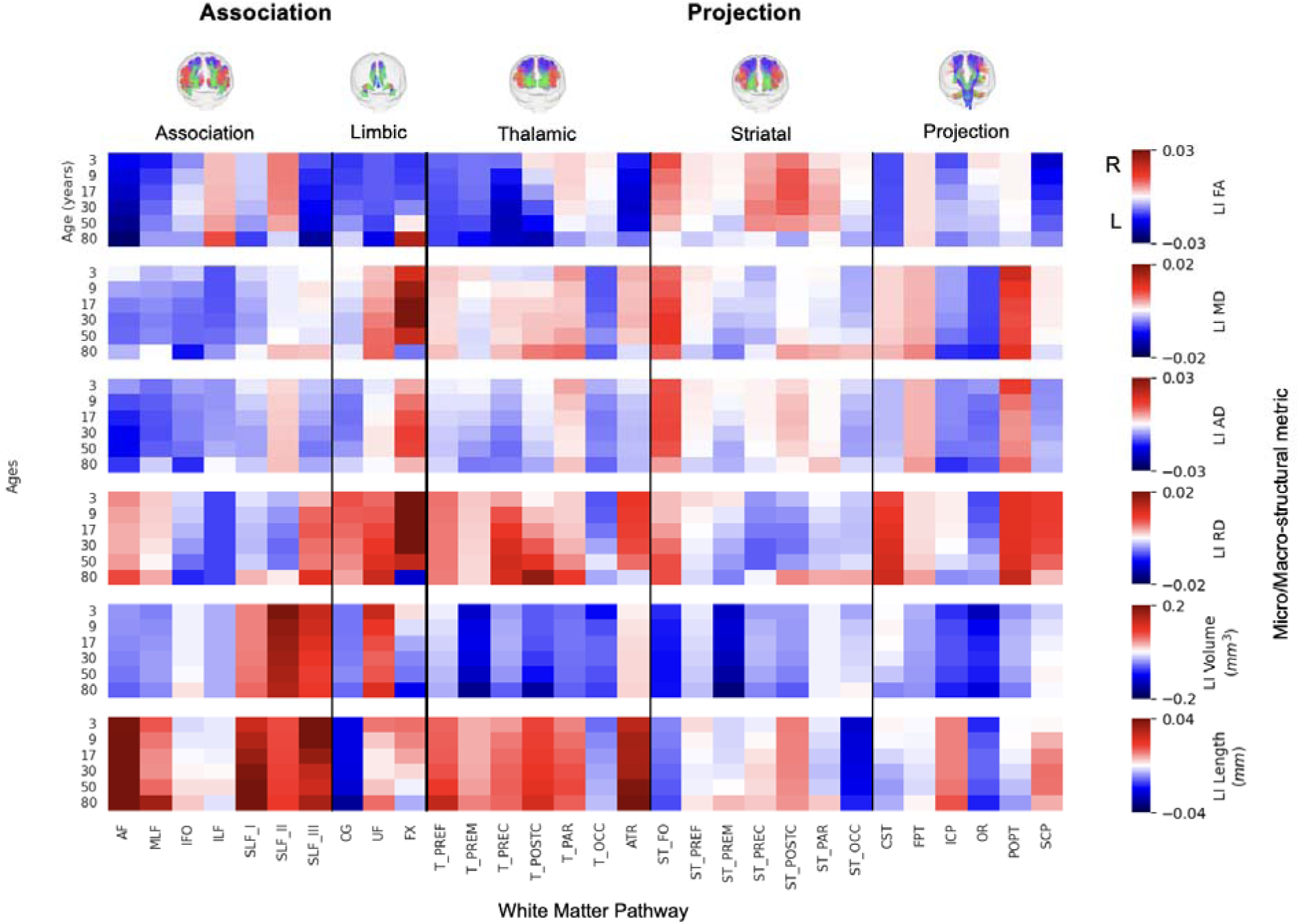
Population-level asymmetry at key lifespan milestones is highly specific to anatomical pathway and structural feature. Heatmaps depict the population-median Lateralization Index (LI) for six key features at six discrete age points (3, 9, 17, 30, 60, and 80 years) across 30 major white matter tracts. The color scale reflects the direction and magnitude of the median asymmetry (red: rightward; blue: leftward), highlighting that distinct patterns of lateralization are already established in early life and continue to evolve across the lifespan.

**Figure 5.**
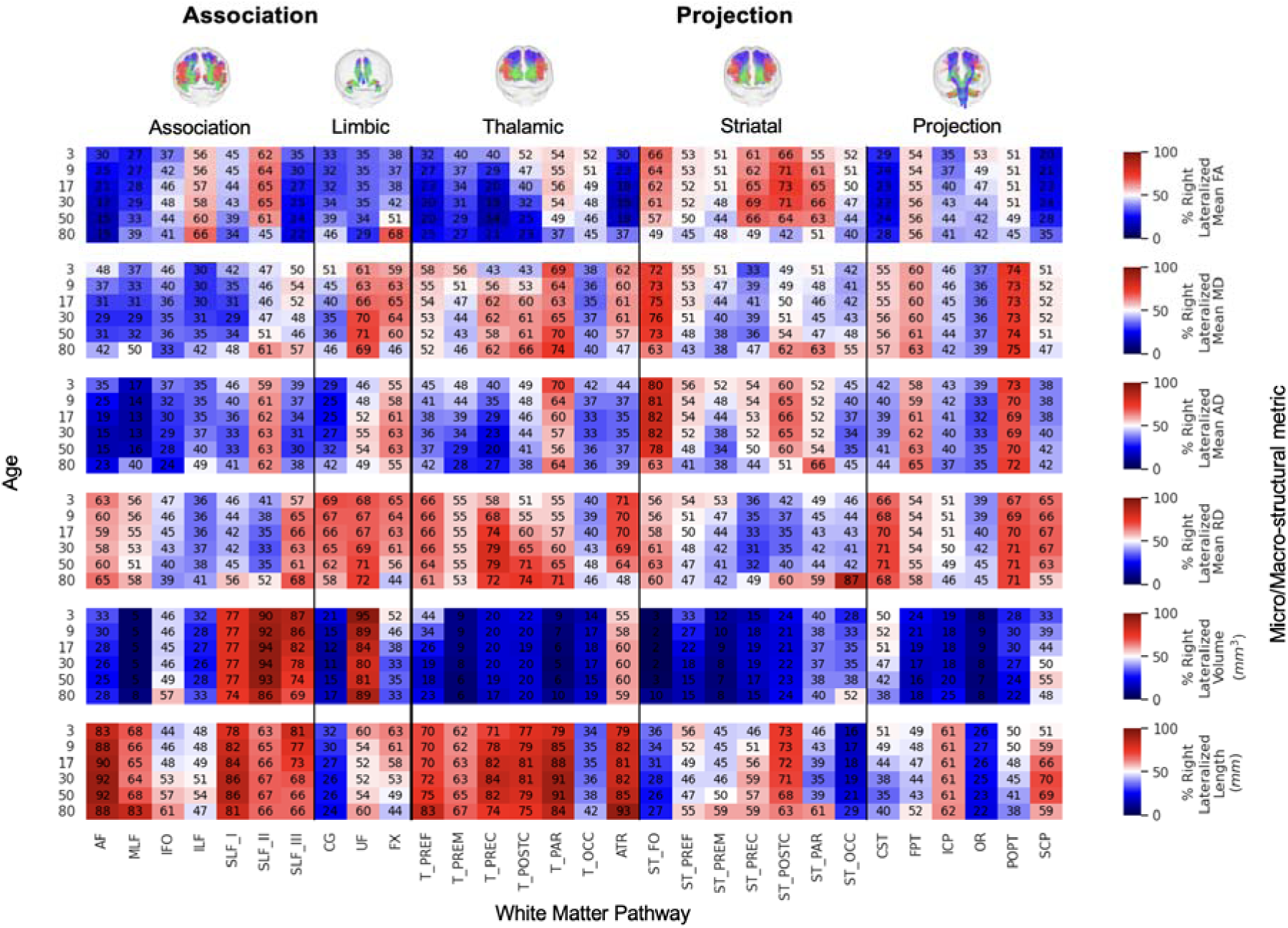
Population prevalence of rightward asymmetry across individuals. This figure complements **Figure 4** by showin the percentage of the population that is right-lateralized for the same features, tracts, and age points. Numerical values and the color scale indicate this prevalence (red: >50% right-lateralized; blue: <50% right-lateralized, i.e., majority left-lateralized). This visualization reveals the consistency of asymmetry across the population, demonstrating that even for tracts with a small median LI, a strong majority of individuals may share the same direction of lateralization.

The results highlight a considerable range in the degree of lateralization across the connectome. Some pathway-feature combinations showed a near-even split in directional preference across the population (i.e., median LI near 0 in **Figure 4** and % right-lateralized near 50% in **Figure 5**). Others exhibited strong and consistent lateralization, with 75-90% or more of individuals showing asymmetry in the same direction (e.g., AF, middle longitudinal fasiculus [MLF], and striato-fronto-orbital [ST_FO] volume). Importantly, many of these distinct patterns were already clearly established in early childhood (3 years old).

Examining trends by feature type revealed broad patterns. For microstructure, FA was typically left-lateralized (negative LIs) across the population for most association and thalamic tracts, with the main exception being the second branch of the fronto-parietal superior longitudinal fasciculus (SLF II), which tended to be right-dominant. In contrast, FA in striatal tracts, was more often right-lateralized in ∼50-70% of the population. Other diffusivity measures (MD and AD) showed a modest but consistent leftward asymmetry across many pathways. For macrostructure, tract volume asymmetries were highly heterogeneous. Strong leftward lateralization (negative LIs) was evident for the AF (∼75% of population), MLF (∼95% of population), ST_FO (>95% of population), optic radiation (OR, >90% of population), and the cingulum bundle (CG; >80%). Strong rightward asymmetry (positive LIs) was observed for all branches (I, II, and III) of the fronto-parietal superior longitudinal fasciculus (SLF_I, 74-77%; SLF_II, 86-95%; and SLF_III, 69-87%), and the uncinate fasciculus (UF, 80-95%). Length also varied considerably; for instance, the length of AF was right-lateralized despite volume being left-lateralized, whereas length of CG was left lateralized alongside its volume. Visual inspection of the heatmaps suggests these patterns are further modulated by age, a dynamic further explored in the final section.

### Do asymmetries reverse direction across the lifespan?

Our analysis of the population median LI trajectories identified instances of “lateralization reversal,” where the typical direction of (population-level) asymmetry shifts from on hemisphere to another across the lifespan (**Figure 6**). This indicates that hemispheric lateralization is not a fixed characteristic but can dynamically evolve with age. As shown in th figure, reversals in the median LI, though relatively sparse, are evident across all major tract groups. There are few patterns that generalize across all pathways or all features, although we note FA tends to reverse from right-to-left (if this occurs), whereas diffusivity measures generally show the opposite left-to-right trend. The timing of these population-level transitions varies considerably across the connectome. For instance, some median LIs reversed direction early in development, while others did so much later in life. This heterogeneity suggests complex and varying degrees of hemispheric plasticity over the lifespan, likely reflecting distinct underlying neurodevelopmental and neurodegenerative processes.

**Figure 6.**
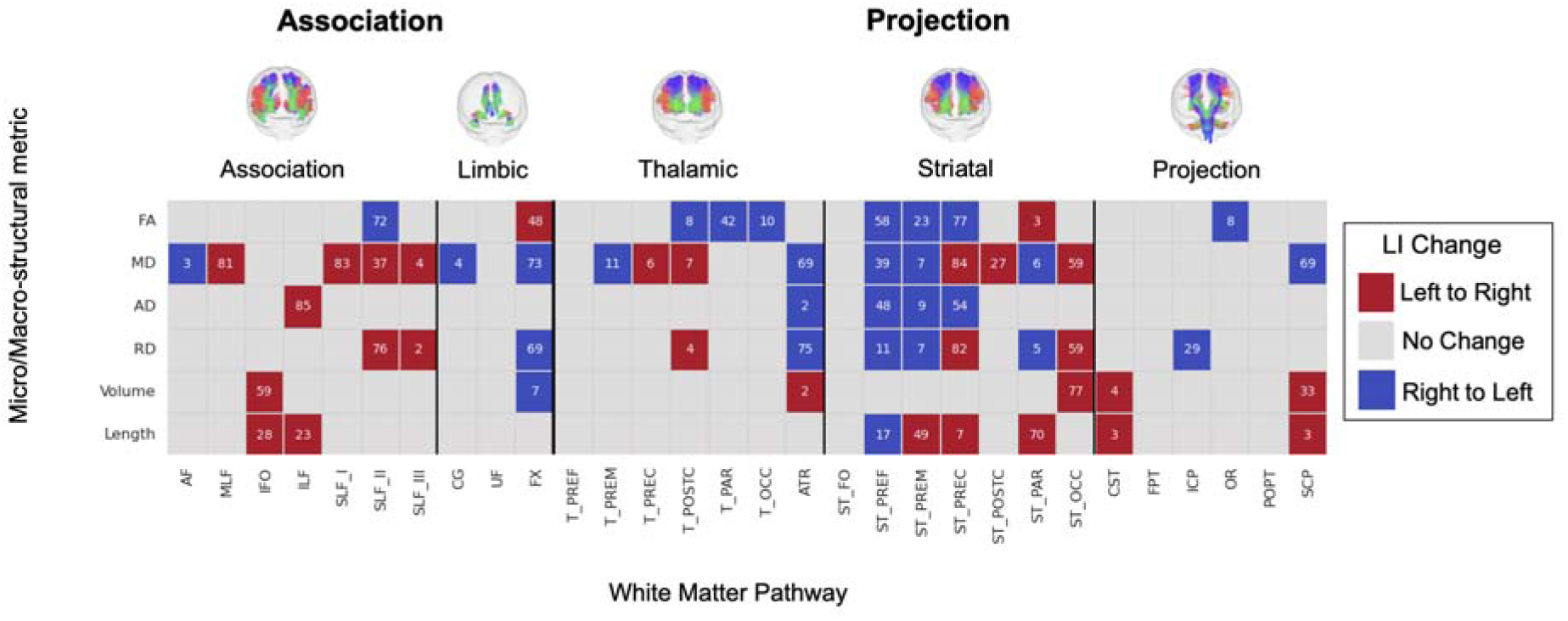
Lifespan trajectories reveal dynamic reversals in the direction of hemispheric asymmetry. Each cell indicates a “lateralization reversal,” where the population-median LI for a given tract-feature combination crosses the zero line. Red cells indicate a transition from leftward to rightward asymmetry (negative to positive LI), while blue cells indicate a transition from rightward to leftward (positive to negative LI). The overlaid numbers represent the age (in years) at which this reversal occurs. The results demonstrate that while reversals are sparse, they occur across all major pathway types and at various life stages.

### How does the magnitude of asymmetry evolve during different life stages?

Finally, we quantified how the magnitude of asymmetry evolves by calculating its rate of change (the slope of the absolute value of the median LI) within six distinct life stages: early childhood (2–5 years), childhood (5–12 years), adolescence (12–20 years), early adulthood (20–40 years), midlife (40–60 years), and older adulthood (60–100 years). **Figure 7** shows these results, where an increasing asymmetry (greater |LI|, meaning *more* leftward or *more* rightward asymmetry) is indicated in red, and decreasing asymmetry (convergence towards symmetry, i.e., *less* leftward or *less* rightward) is shown in blue.

**Figure 7.**
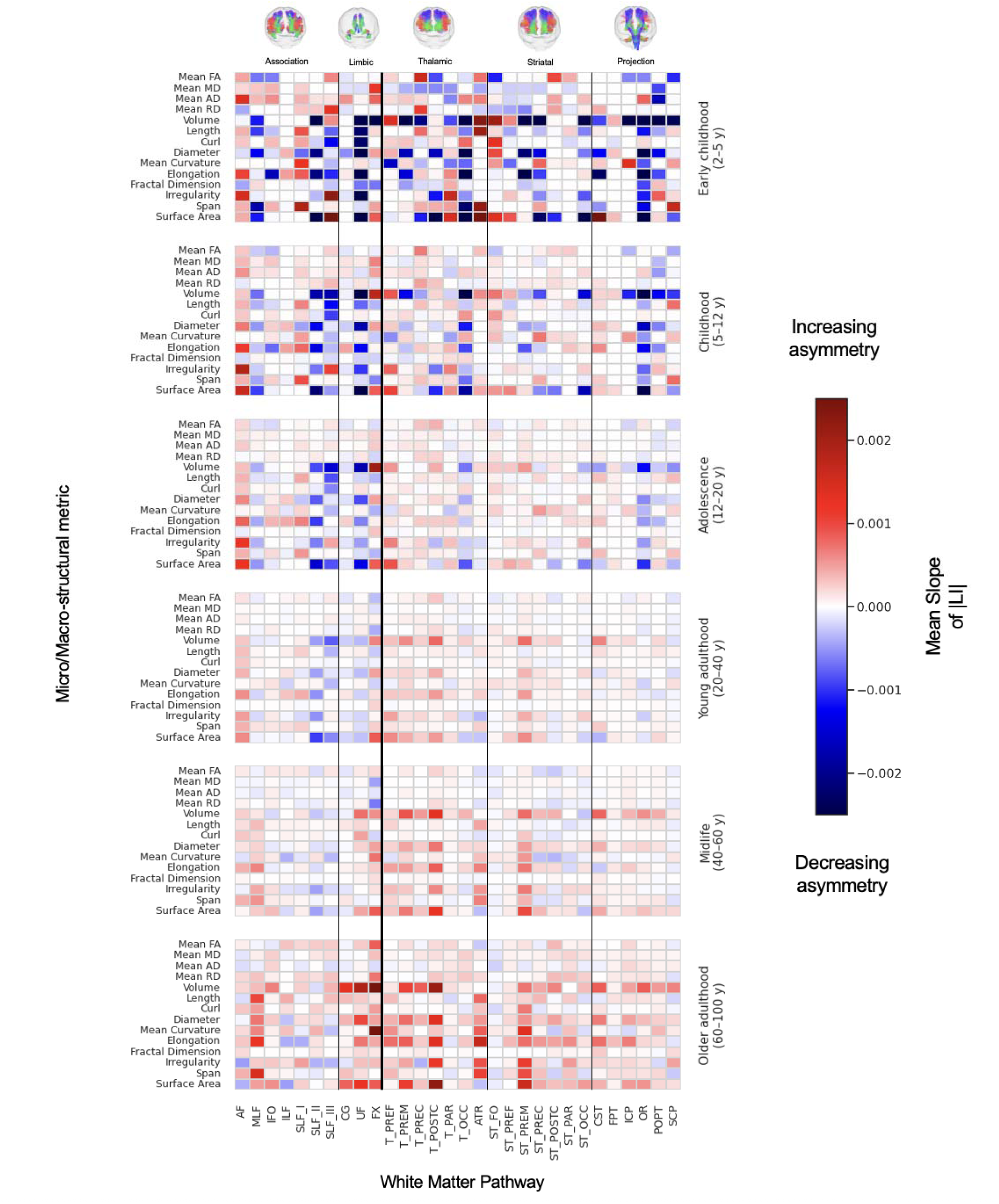
The evolution of asymmetry is a dynamic and age-specific process. This figure shows the mean rate of change (slope) of the population-median absolute lateralization index (|LI|) within four distinct life stages (rows). Each cell represents the slope for a given feature and white matter tract. Warmer colors (red) indicate an increase in the magnitude of asymmetry (strengthening of lateralization), while cooler colors (blue) denote a decrease in the magnitude of asymmetry (attenuation of lateralization). Childhood shows heterogeneous increases and decreases across pathways, adolescence largely continues these trends with smaller slopes, young adulthood is comparatively muted, and later adulthood shows a predominant increase—most marked for macrostructural measures (volume, surface area).

Childhood shows the most heterogenous patterns. Within the same feature class, some pathways show increasing symmetry while other show decreasing. For volume, increasing asymmetry i observed in AF, inferior longitudinal fasciculus (ILF), and several frontal/prefrontal connection (e.g., thalamo-prefrontal [T_PREF], ATR, FPT), whereas SLF-II, SLF-III, UF, and OR tend to decrease. For FA, increases appear in AF, SLF_III, and thalamo-premotor (T_PREM), while ST_FO, MLF, and ILF decrease. This mixture suggests active, pathway-specific refinement in childhood.

Adolescence largely continues the childhood directions but with smaller slopes (i.e., change persist but are less pronounced), consistent with a tapering of developmental remodeling. Young adulthood shows mostly modest shifts, with residual increases in selected macrostructural measures (volume and surface area standing out) and relatively small changes in diffusion metrics.

In middle/late adulthood, a trend emerges where the magnitude of asymmetry generally increases across a wide range of tracts and features. This effect was particularly strong for macrostructural measures like volume and surface area, which had generally been tending towards symmetry in earlier stages of development. For example, the MLF, UF, and nearly all projection pathways showed a strengthening of asymmetry in later life. In contrast, decreases in asymmetry (i.e., tracts becoming more symmetric) were less frequent, though notable exceptions were observed (including the volume of the SLF_II and SLF_III).

## Discussion

This study represents the most comprehensive investigation of white matter asymmetry across the human lifespan. By leveraging a large-scale dataset of over 35,000 individuals from 50 neuroimaging cohorts, we generated normative lifespan trajectories for 6 distinct microstructural and macrostructural features across 30 lateralized long-range white matter pathways. Our principal findings reveal a complex and dynamic brain asymmetry across pathways, features, and the lifespan. First, we demonstrate that white matter asymmetries are widespread, affecting nearly all studied pathways, though the magnitude and lateralization vary considerably. Second, our results show that these asymmetries are feature-dependent, with different or even diverging patterns observed between measures of tissue microstructure and those reflecting pathway macrostructure. Third, we highlight that asymmetry profiles are pathway specific, likely reflecting the unique functional roles and developmental timelines of different neural pathways. Finally, we show that these asymmetries are not static but are highly dynamic, characterized by rapid, heterogeneous changes during childhood and a general trend of becoming more pronounced during the aging process. Together, these lifespan charts quantify the magnitude, direction, and age-specific variability of white-matter asymmetry, providing a reference to interpret individual differences and to study deviations in development, aging, and disease.

### Widespread White Matter Asymmetry and Comparison with Prior Literature

Our study confirms and substantially extends the view of hemispheric asymmetry as a fundamental organizing principle of the brain’s white matter connectome. By charting 30 pathways across the lifespan, we establish that asymmetries are common, yet heterogenous. A key insight from our normative modeling approach is the distinction between the *magnitude* of an asymmetry and its *consistency* across the population. While microstructural asymmetries often had modest population-median LIs, our centile charts reveal that even a small median effect can reflect a strong population consensus, with 70-90% of individuals showing the same directional asymmetry (**Figure 3**). Conversely, a larger median LI did not always guarantee unanimity. This highlights a critical limitation of relying on mean-based analyses alone and underscores the power of a distributional perspective for understanding brain lateralization.

These analyses also highlight that the nature and degree of asymmetry differ across both structural features and anatomical pathways, with many of these distinct lateralization patterns already evident in early infancy. Our lifespan perspective also revealed that inter-individual variability in asymmetry is not uniform (**Figure 3**, centile bands), with the widest centile bands observed in early childhood and in older age. This pattern likely reflects heterogeneous maturational rates and synaptic pruning during development (Dubois et al., 2014; Dubois et al., 2025), and the cumulative impact of diverse genetic, environmental, and health-related factors during aging (Habeck et al., 2017).

Compared with prior reports studying asymmetry - typically smaller, age-restricted, and focused on a few tracts or metrics (**Table 2**, for a review of sample size and age range see **Supplementary Table 2**) - our lifespan curves generally corroborate well-established findings, such as the leftward FA and volumetric asymmetry of the arcuate fasciculus in adulthood, validating our large-scale harmonization. However, our data also help clarify previously conflicting reports in the literature (Dulyan et al., 2025) - where some studies demonstrated structural features to be left-lateralized and others demonstrated those same features to be right-lateralized - particularly for association pathways like the inferior fronto-occipital fasciculus (IFO), and UF. For the IFO, some studies report volume to be left-lateralized (H. W. Powell et al., 2006), whereas others (Wu, Sun, & Wang, 2016) (and our results) suggest there is no lateralization of IFO volume. Similarly, for the UF, some studies have demonstrated FA to be right-lateralized (Parekh et al., 2023), whereas others (Hasan et al., 2009; H. W. Powell et al., 2006) (and our results) indicate leftward lateralization. For the CST, some studies report left-lateralized microstructure (Angstmann et al., 2016; Demnitz et al., 2021; Parekh et al., 2023), whereas others (Parekh et al., 2023; Schmithorst, Holland, & Dardzinski, 2008) (and our results) indicate right-lateralized microstructure with left-lateralized macrostructure - differences that likely reflect metric choice and age window.

**Table 2.**
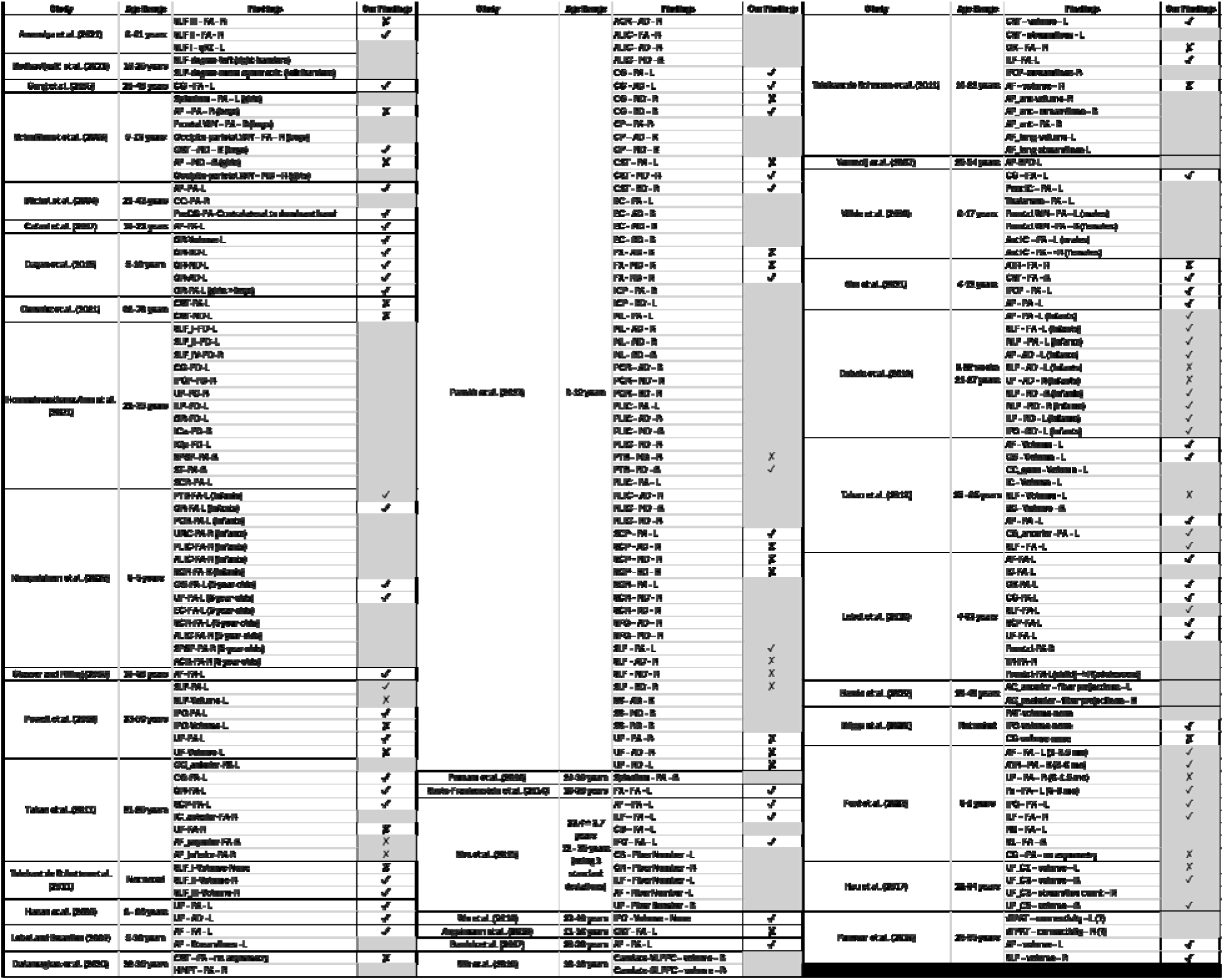
Comparison of our findings and those previously demonstrated in the literature. IZ = agreement between our findings and those presented in the literature. X = disagreement between our findings and those presented in the literature. * (in “Findings”) = tract or measure not measured in our work. * (in “Findings”) + IZ/X = tract represented in our data as conglomerate or as multiple tracts, though still allowing for gross comparison. In cases where tract abbreviations presented in the literature differed from those of our tracts, we presented the tract name using our abbreviations.

Several factors explain discrepancies and the additional patterns detected here. First is scale: with N = 35,120, our centile modelling estimates population trajectories rather than relying on small-sample inference. This makes our study particularly powerful compared to smaller studies and enables its future use as a reference for lifespan asymmetry in healthy patients. Indeed, a major motivation for using a GAMLSS-based normative framework was its ability to estimate age-varying distributions while explicitly accounting for between-cohort heterogeneity (e.g., scanner/protocol differences) within a single model (Rigby & Stasinopoulos, 2005). Second, continuous age modelling (rather than single time points or assumption of homogeneity across age) exposes direction changes and non-linear trends that limited-age designs miss. Third, tract definition may vary across studies (K. G. SchillingF. Rheault, et al., 2021). Automated, standardized segmentation (e.g., TractSeg) (Wasserthal et al., 2018) reduces protocol variance but will not replicate every manual or atlas-based definition used previously. Finally, is the choice of features, as micro-and macrostructural features measure different aspects of pathway biology and can often diverge. By providing comprehensive, openly available lifespan asymmetry charts, our work offers a stable reference point to help reconcile these inconsistencies and guide future research.

We additionally generated handedness-specific lifespan charts in the subset of cohorts with handedness information available (12 cohorts; *N* = 14,220; analyses focused on right-vs left-handed individuals). Across exemplar tracts and features, handedness-related differences in the modeled median LI were consistently small (Cohen’s *d* [calculated at every integer age 1-80 and averaged across the lifespan] typically < 0.08 and often < 0.02; **Supplementary Figure 1**), indicating that handedness explains little additional variance in tract-level asymmetry in these data. This analysis is necessarily more weakly constrained than our primary models (i.e., does not strictly align with **Figure 3**) because handedness was unavailable for most cohorts and the left-handed subgroup was smaller and not evenly distributed across the lifespan - factors known to reduce precision when estimating centile curves within subgroups (i.e., optimally designed growth reference studies often require very large samples per group (Cole, 2021)). Despite this, the minimal handedness effects observed here are broadly consistent with prior reports that structural asymmetry relates only weakly to handedness (J. L. Powell et al., 2012) (Groen, Whitehouse, Badcock, & Bishop, 2013) (Guadalupe et al., 2014; Kong et al., 2018), even as functional lateralization and language-network connectivity can differ by handedness (Wiberg et al., 2019).

### Divergent Asymmetries of Microstructural and Macrostructural Features

Our findings demonstrate that white matter asymmetry is not “all encompassing” within a given pathway. Rather, it is highly feature dependent, meaning a tract broadly described as “left-lateralized” may not show leftward “dominance” for all its micro-and macrostructural characteristics. For instance, a pathway may exhibit leftward asymmetry for FA and other diffusivities (MD/AD) yet display rightward asymmetry for RD, as is seen in the arcuate fasciculus (**Figure 3**); or show leftward asymmetry in FA while its overall volume is greater in the right. These examples are widespread throughout our investigation (**Figures 4, 5**). This divergence highlights why simplistic labels such as “dominant” hemisphere for a tract or generalized statements about greater “white matter integrity” can be insufficient or even misleading without specifying which feature (reflecting distinct biological properties) is lateralized. Indeed, diffusion tensor metrics like FA and RD are sensitive to processes like myelination and axonal packing, AD to axonal structure and coherence, and MD to overall water diffusivity, each providing unique insights into pathways’ structure (Beaulieu, 2002). The field must continue to investigate the biological and functional significance of these feature-specific asymmetries to better understand brain lateralization.

This divergence is biologically plausible and offers insight into brain organization on multiple scales. Macrostructural asymmetries could represent an earlier, more genetically-programmed aspect of development that establishes the gross architecture of a pathway (Budisavljevic, Castiello, & Begliomini, 2021; Habeck et al., 2017). In contrast, microstructural asymmetries might reflect more dynamic, experience-dependent processes, such as the activity-dependent myelination that refines and optimizes a circuit for a specific function (Groen et al., 2013; J. L. Powell et al., 2012). This dissociation underscores why simplistic labels like a “dominant” hemisphere or generalized statements about “white matter integrity” can be misleading (Kong et al., 2018). Interpreting lateralization and its functional or dysfunctional implications requires considering both large-scale architecture and cellular-level microstructure of white matter pathways.

Diffusion tensor indices (FA, MD, AD, RD) should not be interpreted as independent biological measures, as they are different summaries of the same tensor model (Wheeler-Kingshott & Cercignani, 2009). In particular, FA is mathematically determined by the tensor eigenvalues and therefore depends on the relative behavior of axial and radial diffusivity. As a result, cases where FA laterality differs from RD (or AD) laterality should be interpreted as complementary descriptors of how diffusion parallel versus perpendicular to the principal fiber direction differs between hemispheres, rather than as evidence for unrelated or competing biological processes.

At the same time, diffusion-derived metrics are sensitive but not specific to myelin or any single microstructural substrate; FA for example, reflects not only myelin but also broader features of tissue microstructure and organization (e.g., membrane/axon packing, caliber, and fibre geometry) (Beaulieu, 2002), and - like all tensor-derived indices - its interpretation can be biased in voxels with complex fibre architecture (e.g., crossing/branching bundles) and partial-volume with other tissues (Pierpaoli, Jezzard, Basser, Barnett, & Di Chiro, 1996). For this reason, diffusion-based laterality is usefully interpreted alongside complementary contrasts that more directly index myelination. As an example, myelin water fraction imaging provides a more myelin-specific estimate with strong histological correspondence and has also demonstrated hemispheric asymmetries in healthy participants and early development (Deoni, Dean, O’Muircheartaigh, Dirks, & Jerskey, 2012), including associations with behavior (reading, language) (Beaulieu et al., 2020; O’Muircheartaigh et al., 2013) and genetics (Ocklenburg et al., 2019). Together, these findings support the view that a subset of the lifespan asymmetry patterns observed here - particularly those involving diffusivity-based measures - may partly reflect hemispheric differences in myelination, while motivating future multimodal studies that directly combine diffusion modeling with quantitative myelin imaging or biophysical modeling (Novikov, Fieremans, Jespersen, & Kiselev, 2018).

### Pathway-Specific Patterns of White Matter Asymmetry

While previous work has generated such charts for grey matter (Bethlehem, Seidlitz, et al., 2022) and white matter (Michael E. Kim et al., 2025), this is the first study to specifically chart tract-level asymmetry by creating population-level curves (describing the population-distribution) across age. From a basic neuroscience perspective, this resource helps quantify how neural circuits differ between hemispheres, supporting ongoing efforts to relate large-scale neuroanatomical organization to functional specialization (Forkel, Friedrich, Thiebaut de Schotten, & Howells, 2022).

Across pathways, we found that asymmetry profiles are tract-specific and align with several recurring themes described in prior literature. For example, as noted previously, the arcuate fasciculus shows a robust leftward bias for several features across much of the lifespan, consistent with the tract’s central role in the dorsal language system. In contrast, frontoparietal association pathways (most notably SLF I,II,III) show strong rightward macrostructural asymmetry in a large majority (70-85%) of individuals (**Figure 5**), a pattern that has been linked to right-hemisphere dominant attentional/visuospatial networks (Dulyan et al., 2025; Thiebaut de Schotten, Dell’Acqua, et al., 2011), and consistent with right-lateralized IFO volume that has been interpreted as a structural basis for the right hemisphere’s strengths in visuospatial integration (Thiebaut de Schotten, Ffytche, et al., 2011). Together, these examples (and the widespread asymmetries across all pathways studied) highlight that structural lateralization is not limited to a single “language tract,” but is a distributed property of multiple functional systems. These pathway-specific reference distributions can help guide future work aiming to link particular asymmetry profiles to individual differences in cognition, behavior, and vulnerability to neurological or psychiatric disorders.

A key point for interpretation is that tract asymmetries often co-occur with well-established cortical asymmetries in the same functional systems (Good et al., 2001; Minkova et al., 2017; Ocklenburg, Friedrich, Güntürkün, & Genç, 2016; Toga & Thompson, 2003), suggesting coordinated hemispheric specialization across gray-matter nodes and the long-range pathways that connect them. In the language network, Roll (2024) reports pronounced leftward surface-area asymmetry in Heschl’s gyrus and a dissociation within the inferior frontal gyrus, where pars opercularis shows leftward surface-area asymmetry while anterior IFG subdivisions show the opposite pattern (Roll, 2024). Roll also highlights a broader anatomical signature in left-hemisphere language regions - larger surface areas paired with thinner cortex and a higher white-to-gray matter ratio **–** a profile consistent with a more myelinated and efficient substrate for rapid categorical processing. These cortical asymmetries provide an anatomical context for the left-lateralized AF that we observe: the AF is the major long-range association pathway linking posterior temporal/auditory regions with posterior inferior frontal speech regions, and multiple studies have shown that inter-individual differences in arcuate laterality relate to functional language lateralization. In our data, this same circuit shows complementary tract-level asymmetries across much of the lifespan, including leftward asymmetry in AF volume and FA (with a rightward asymmetry in mean streamline length), consistent with hemispheric differences in both pathway architecture and diffusion-derived tissue organization. These gray-and white-matter asymmetries likely reflect partly shared developmental and genetic influences, and our lifespan charts provide a starting point for testing how they covary with functional lateralization and behavior across age.

Beyond association pathways, our results reveal that thalamic projection tracts tend to exhibit leftward microstructural asymmetry while striatal projection tracts show the opposite pattern, despite their partial anatomical overlap. This divergence is consistent with prior work demonstrating lateralized, circuit-specific differences in thalamocortical versus corticostriatal circuitry and likely reflects fundamental organizational differences in their connectivity and functional roles (Foster et al., 2021; Martel & Galvan, 2022; Shepherd & Yamawaki, 2021). Thalamocortical pathways demonstrate reciprocal and nonreciprocal connections with layer-specific cortical termination patterns that support hierarchical information flow and sensory integration, whereas corticostriatal pathways exhibit greater convergence and divergence to enable integration of distributed cortical inputs for action selection and motor–cognitive control (Martel & Galvan, 2022; Shepherd & Yamawaki, 2021; Wolff, Ko, & Ölveczky, 2022).

### Dynamic Nature of Asymmetry Across the Lifespan

A central finding of this work is that white matter asymmetries are not static but follow dynamic trajectories across the lifespan (**Figure 3**). These dynamics are evident in both the population median LI trajectories and the corresponding population variability. While the typical magnitude of the median LI for many features, particularly microstructural ones, often remains within a relatively constrained range (e.g., frequently staying below +/-0.1), especially during most of adult life, their specific values and trajectories show population-level changes over decades. This evolution varies considerably by pathway: some tracts exhibit clear shifts and modulations in their median asymmetry profiles across developmental and aging periods (e.g. which tracts), while others maintain a more consistent pattern of lateralization and population distribution throughout much of life (e.g. AF). The periods of most pronounced change in these asymmetry trajectories - reflecting significant neural reorganization generally occur during very early development and again in later life during healthy aging. It is important to highlight the cross-sectional nature of the current dataset; because of this, we cannot (or do not) follow individual “change” but rather cross-sectional median and ranges across the population.

Interpreting lifespan trajectories of the laterality index (LI) requires considering the underlying bilateral trajectories from which LI is computed. Because LI is a normalized contrast of left-and right-hemisphere tract properties, the same apparent change in LI (e.g., attenuation or strengthening) can arise through multiple mechanisms: a decrease in the feature on the initially dominant side, an increase on the contralateral side, or differences in the timing and slope of change across hemispheres. This is illustrated in Figure 8, which juxtaposes exemplar left-and right-hemisphere lifespan curves for tract microstructure with the corresponding LI trajectory. Large, stable LIs occur when the two hemispheric trajectories remain separated across age (e.g., AF), whereas small LIs occur when trajectories track closely (e.g., CST). In other cases, LI shifts or reversals can emerge when left and right pathways evolve at different rates such that the trajectories converge, diverge, or cross (e.g., FX). Together, this reinforces that white-matter asymmetry is an emergent property of paired hemispheric development and aging, rather than a change attributable to one hemisphere alone.

**Figure 8.**
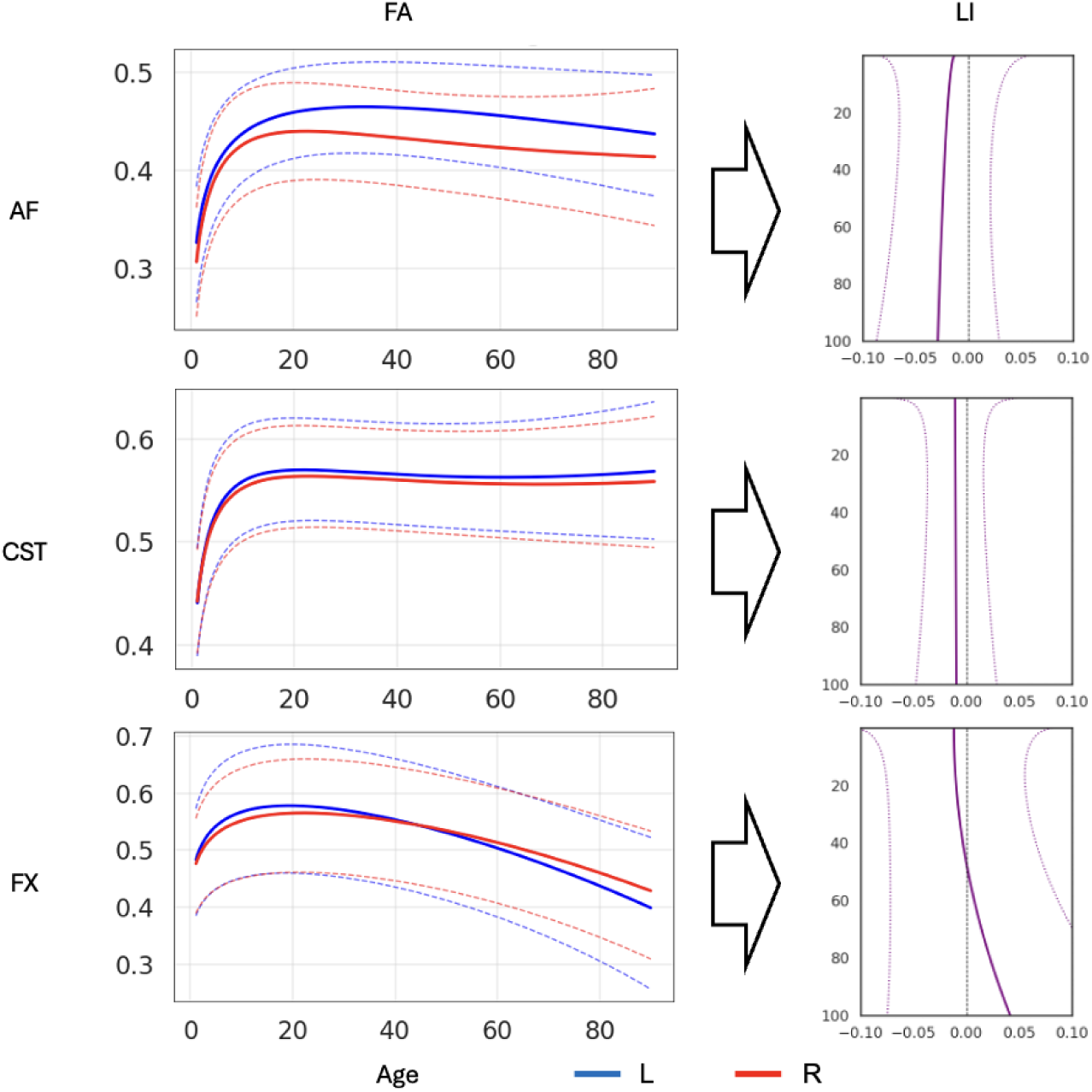
Interpreting LI trajectories in the context of bilateral lifespan trajectories. Left-and right-hemisphere lifespan trajectories for exemplar tract–feature combinations (left panels) are shown alongside the corresponding LI trajectories (right panel). The figure illustrates that larger asymmetries arise when left and right trajectories remain widely separated, smaller asymmetries arise when trajectories are closely aligned, and LI shifts or reversals can occur when hemispheric trajectories change at different rates across age. Importantly, LI curves were estimated by computing LI at the individual level and fitting normative models to those LI values; they are not derived by algebraically differencing the plotted left/right population trajectories.

This interpretation is consistent with prior diffusion MRI lifespan studies (Beck et al., 2021; Michael E. Kim et al., 2025; Lebel et al., 2012; Schilling et al., 2023; Villalón-Reina et al., 2024; A. H. Zhu et al., 2024) showing that white-matter microstructure follows tract-specific, often non-linear developmental and aging trajectories rather than a single uniform pattern across contralateral pathways or different pathways.

We also demonstrate, for the first time to our knowledge, the presence of “lateralization reversal”, where the typical direction of asymmetry for a given feature shifts across the lifespan. This phenomenon occurred in almost every tract we measured and for at least one, and often more, measures of connectivity (**Figure 6**), although the overall frequency was indeed spars across pathways and features. While previous studies have demonstrated patterns of development of asymmetry over short age ranges (Ford, Ammar, Li, & Shultz, 2023; Venla Kumpulainen, 2024; Lebel, Treit, & Beaulieu, 2019), identifying specific reversal points is possible by including data across the entire lifespan. The neurodevelopmental and clinical significance of this phenomenon is, to our knowledge, completely unexplored.

Finally, our analysis reveals another novel lifespan trend: the degree of white matter asymmetry shows a general tendency to increase from mid-life to old age (**Figure 7**). While asymmetry in infancy/development has been a focus of prior research (Dubois et al., 2016), its evolution during aging is less understood with limited prior investigations (Demnitz et al., 2021). Our results demonstrate that several features (volume, diameter, elongation, irregularity, and surface area) become more asymmetric with age across most pathways, and many other features show similar trends. We hypothesize that these findings are reflective of natural age-related neurodegenerative processes – occurring in individuals over 50 years old – where one hemisphere’s pathways (or specific aspects of their structure) are more vulnerable or decline at a faster rate than their contralateral counterpart, leading to the amplifying initial subtle asymmetries. Future studies are crucial to determine the neurocognitive and clinical impact, if any, of these age-related increases in white matter asymmetry.

### Functional Implications and Clinical Relevance of White Matter Asymmetry

Our findings also carry potential clinical implications. As research accumulates, white matter asymmetry measures could serve as biomarkers for certain conditions such as schizophrenia (Miyata et al., 2012) and Parkinson’s disease (Y. Zhu et al., 2023) or as indices of typical vs. atypical brain development, such as in autism spectrum disorder (Joseph et al., 2014). Asymmetry is a particularly attractive biomarker since it is inherently normalized to the individual via its computation as ratios or differences within the same scan. In children with neurodevelopmental disorders, it has been demonstrated that lateralization indices of diffusion metrics might serve as clinically useful imaging biomarkers, helping to detect sensory processing dysfunction with less confounding variance than absolute measures (Parekh et al., 2023). In neurodegenerative diseases, asymmetry often manifests as an asymmetrical onset of pathology. Parkinson’s disease typically begins with unilateral motor deficits (reflecting greater degeneration of one side’s nigrostriatal pathway), and patients correspondingly show asymmetric cortical changes in several hemispheric regions (Claassen et al., 2016); a similar investigation into asymmetrical changes in white matter structural connectivity has not yet been conducted. With continued development, detection of such asymmetry measures can complement other markers to improve early detection of atypical brain development or pathology.

The role of asymmetry across disease states has only begun to be characterized. When considering clinical populations, white matter asymmetry has been a topic of intense investigation. Research in autism, for example, has observed that typical lateralization patterns are often reduced or altered. Children with autism spectrum disorer have been shown to have significantly diminished asymmetry of white matter microstructure, with many of the normal left-right differences in FA, AD, and RD being smaller in magnitude in autism spectrum disorder, indicating a more symmetrical white matter organization (Carper et al., 2016; Mundorf et al., 2021). Schizophrenia is another condition where white matter asymmetry is studied: some patients show a reduction or reversal of asymmetry in the AF and SLF, among others, with reduced asymmetry in the FA being seen in patients experiencing auditory hallucinations (Catani et al., 2011), possibly underlying the disturbed lateralization of language processing in schizophrenia (Mundorf et al., 2021). In such investigations of pathology-associated asymmetries, our work can serve as a cornerstone for future comparisons. By providing a multitude of tracts and measures modeled across the lifespan, our charts can reasonably be used as healthy controls in any study which seeks to investigate a particular tract, connectivity measure, and/or critical age range in an abnormal brain state. Our findings in healthy controls, therefore, potentially enables the discovery of different asymmetries that arise in pathological states, providing insight into the mechanism of a given disease.

## Limitations

Our study had notable limitations. We note the results for the fornix (FX) should be interpreted with caution as they could be a result of technical tractography artifacts rather than a biological changes; while this is particularly true of FX, such considerations can be grossly applied to all tractography methods. Macrostructural features are susceptible to the methods used to grid data and so can vary between studies. Similarly, different protocols for delineation of pathways can lead to different regions being included or excluded from a given pathway, lending further complexity to the matter of comparing study results. Lastly, there is overlap in the regions which define given pathways, meaning that microstructural measures in those regions are not unique to the given pathway. This is an inherent limitation of tract-based summaries of voxel-wise diffusion tensor metrics: even when measures are weighted by streamline density or restricted to a tract mask, the underlying scalar values are still estimated at the voxel level and can reflect signal contributions from other pathways in regions of crossing/overlap (K. G. Schilling, C. M. W. Tax, et al., 2021).

## Conclusion

This study provides the most extensive characterization of white matter asymmetry across the human lifespan to date, generating comprehensive normative trajectories for 6 structural features across 30 distinct white matter pathways in over 35,000 healthy individuals from infancy to old age. Our key findings demonstrate that white matter asymmetries are widespread yet highly specific to pathways and structural features. Furthermore, these asymmetries are not static but exhibit dynamic changes throughout development and aging, including reversals in direction and a general trend of increasing asymmetry magnitude in later life. These lifespan charts of white matter asymmetry offer a resource for advancing our understanding of brain lateralization principles, investigating inter-individual variability in brain structure, and providing a reference to assess deviations in neurodevelopmental, aging, and clinical populations.

## Data Availability

Population curves for each pathway and each feature are in format npy and available on Zenodo [https://doi.org/10.5281/zenodo.18202413]. Code for GAMLLS fitting is at https://github.com/MASILab/Asymmetry/src/fit_model_pynm.py and https://github.com/MASILab/Asymmetry/src/fit_model_handedness_pynm.py. Code to generate figures is at https://github.com/MASILab/Asymmetry/src/main_text_figures.ipynb, https://github.com/MASILab/Asymmetry/src/handedness_figures.ipynb, and https://github.com/MASILab/Asymmetry/src/dataset_plots.ipynb. Additionally, the GAMLLS models for each pathway and each feature are available on Zenodo.

## Declaration of Generative AI and AI-assisted Technologies in the Writing Process

During the preparation of this work the authors used ChatGPT, an AI language model developed by OpenAI, and Gemini, an AI language model developed by Google, in order to assist in rephrasing the text in this paper for clarification. After using this tool, the authors reviewed and edited the content as needed and takes full responsibility for the content of the publication.

## Acknowledgements and Funding

This work was supported in part by the National Institute of Health through NIH awards K01-EB032898 (Schilling) and K01-AG073584 (Archer), grant number 1R01EB017230-01A1 (Landman), K24-AG046373 (Jefferson), and ViSE/VICTR VR3029, UL1-TR000445, and UL1-TR002243. This work was supported by NIA grants R01-AG034962 (Jefferson), R01-AG056534 (Jefferson), R01-AG062826, and Alzheimer’s Association IIRG-08-88733 (Jefferson) and the NICHD, R01-HD114489 (Vinci-Booher). This work was supported by the Alzheimer’s Disease Sequencing Project Phenotype Harmonization Consortium (ADSP-PHC) that is funded by NIA (U24 AG074855, U01 AG068057 and R01 AG059716). This work was conducted in part using the resources of the Advanced Computing Center for Research and Education (ACCRE) at Vanderbilt University, Nashville, TN. We appreciate the National Institute of HealthS10 Shared Instrumentation grant 1S10OD020154-01, and grant 1S10OD023680-01 (Vanderbilt’s High-Performance Computer Cluster for Biomedical Research). This work was also supported in part by Intramural Research Program of the National Institute on Aging, NIH.

Research reported in this publication was supported by NIGMS of the National Institutes of Health under award number T32GM007347 and T32GM152284.

This work was supported by the Donders Mohrmann Fellowship on ‘Neurovariability’ No. 2401515 (Stephanie J. Forkel) and the Dutch Research Council NWO Aspasia Grant ‘Human individuality: phenotypes, cognition, and brain disorders’ (Stephanie J. Forkel).

Data and/or research tools used in the preparation of this manuscript were obtained from the National Institute of Mental Health (NIMH) Data Archive (NDA). NDA is a collaborative informatics system created by the National Institutes of Health to provide a national resource to support and accelerate research in mental health. This manuscript reflects the views of the authors and may not reflect the opinions or views of the NIH or of the Submitters submitting original data to NDA.

This research was supported by the Intramural Research Program of the NIH, National Institute on Aging. The contributions of the NIH authors were made as part of their official duties as NIH federal employees, are in compliance with agency policy requirements, and are considered Works of the United States Government. However, the findings and conclusions presented in this paper are those of the authors and do not necessarily reflect the views of the NIH or the U.S. Department of Health and Human Services.

We acknowledge the data provided by several initiatives:

HABS-HD: Research reported on this publication was supported by the National Institute on Aging of the National Institutes of Health under Award Numbers R01AG054073, R01AG058533, R01AG070862, P41EB015922 and U19AG078109. The content is solely the responsibility of the authors and does not necessarily represent the official views of the National Institutes of Health.

PING: The Pediatric Imaging, Neurocognition, and Genetics (PING) dataset was collected and released openly to contribute to the assessment of typical brain development in a pediatric sample (RC2DA029475-01), (https://www.sciencedirect.com/science/article/pii/S1053811915003572).

HCP: The data used in this study come from the Human Connectome Project, which aims to map the structural connections and circuits of the brain and their relationships to behavior by acquiring high-quality magnetic resonance images. We used diffusion MRI data from the Baby Connectome Project (HCPBaby), the Human Connectome Project Development (HCPD) study, the Human Connectome Project Young Adult (HCP) study, and the Human Connectome Project Aging (HCPA) study.

Research data from the Infant Brain Imaging Study (IBIS) dataset come from the IBIS Autism project, a collaborative effort by investigators to conduct a longitudinal MRI/DTI and behavioral study of infants at high risk for autism (R01HD055741-01).

NKI-RS: The Enhanced NKI-RS is a large cross-sectional sample of brain development, maturation and aging, that is currently funded by the NIMH (BRAINS R01MH094639-01; PI Milham) and Child Mind Institute (PI Milham) to characterize 1000 community-ascertained participants using state-of-the-art multiplex imaging-based resting state fMRI (R-fMRI) and diffusion tensor imaging (DTI), genetics, and a broad neurobehavioral phenotypic characterization protocol. Data were acquired from the DSI studio website: (https://brain.labsolver.org/nki_rockland.html).

HBN: The Healthy Brain Network (HBN) is an ongoing initiative focused on building a biobank of data from 10,000 children and adolescents (ages 5-21) in the New York City area (https://www.nature.com/articles/sdata2017181). Data were acquired from the DSI studio website: (https://brain.labsolver.org/hbn.html).

Humphreys: This work was supported by the Jacobs Foundation Early Career Research Fellowship (2017-1261-05) (Humphreys); National Science Foundation CAREER Award (2042285) (Humphreys); Brain and Behavior Research Foundation John and Polly Sparks Foundation Investigator Award (29593) (Humphreys); Vanderbilt Institute for Clinical and Translational Research Grant (VR53419) (Humphreys); Vanderbilt Strong Grant; Vanderbilt Kennedy Center Grant (Humphreys); National Institute of Mental Health (R01MH129634) (Humphreys).

ADNI: Data used in the preparation of this article were obtained from the Alzheimer’s Disease Neuroimaging Initiative (ADNI) database (adni.loni.usc.edu). The ADNI was launched in 2003 as a public-private partnership, led by Principal Investigator Michael W. Weiner, MD. The primary goal of ADNI has been to test whether serial magnetic resonance imaging (MRI), positron emission tomography (PET), other biological markers, and clinical and neuropsychological assessment can be combined to measure the progression of mild cognitive impairment (MCI) and early Alzheimer’s disease (AD).

Data collection and sharing for ADNI were supported by National Institutes of Health Grant U01-AG024904 and Department of Defense (award number W81XWH-12-2-0012). ADNI is also funded by the National Institute on Aging, the National Institute of Biomedical Imaging and Bioengineering, and through generous contributions from the following: AbbVie, Alzheimer’s Association; Alzheimer’s Drug Discovery Foundation; Araclon Biotech; BioClinica, Inc.; Biogen; Bristol-Myers Squibb Company; CereSpir, Inc.; Cogstate; Eisai Inc.; Elan Pharmaceuticals, Inc.; Eli Lilly and Company; EuroImmun; F. Hoffmann-La Roche Ltd and its affiliated company Genentech, Inc.; Fujirebio; GE Healthcare; IXICO Ltd.; Janssen Alzheimer Immunotherapy Research & Development, LLC.; Johnson & Johnson Pharmaceutical Research & Development LLC.; Lumosity; Lundbeck; Merck & Co., Inc.; Meso Scale Diagnostics, LLC.; NeuroRx Research; Neurotrack Technologies; Novartis Pharmaceuticals Corporation; Pfizer Inc.; Piramal Imaging; Servier; Takeda Pharmaceutical Company; and Transition Therapeutics. The Canadian Institutes of Health Research is providing funds to support ADNI clinical sites in Canada. Private sector contributions are facilitated by the Foundation for the National Institutes of Health (www.fnih.org). The grantee organization is the Northern California Institute for Research and Education, and the study is coordinated by the Alzheimer’s Therapeutic Research Institute at the University of Southern California. ADNI data are disseminated by the Laboratory for Neuro Imaging at the University of Southern California.

VMAP: Study data were obtained from the Vanderbilt Memory and Aging Project (VMAP). VMAP data were collected by Vanderbilt Memory and Alzheimer’s Center Investigators at Vanderbilt University Medical Center. This work was supported by NIA grants R01-AG034962 (PI: Jefferson), R01-AG056534 (PI: Jefferson), R01-AG062826 (PI: Gifford), and Alzheimer’s Association IIRG-08-88733 (PI: Jefferson).

Data/Processes/Plans/Concepts (select as appropriate) used in the preparation of this article were obtained from the Healthy Brain and Child Development (HBCD) Study (Home - Healthy Brain and Child Development). This is a multisite, longitudinal study designed to recruit approximately 7,500 families and follow them from pregnancy to early childhood. The HBCD Study is supported by the National Institutes of Health and additional federal partners under award numbers U01DA055352, U01DA055353, U01DA055366, U01DA055365, U01DA055362, U01DA055342, U01DA055360, U01DA055350, U01DA055338, U01DA055355, U01DA055363, U01DA055349, U01DA055361, U01DA055316, U01DA055344, U01DA055322, U01DA055369, U01DA055358, U01DA055371, U01DA055359, U01DA055354, U01DA055370, U01DA055347, U01DA055357, U01DA055367, U24DA055325, U24DA055330. A full list of supporters is available at https://hbcdstudy.org/about/federal-partners/. A listing of participating sites and a complete listing of the study investigators can be found at Recruitment Sites - Healthy Brain and Child Development. HBCD consortium investigators designed and implemented the study and/or provided data but did not necessarily participate in the analysis or writing of this report. This manuscript reflects the views of the authors and may not reflect the opinions or views of the NIH or HBCD consortium investigators.

BLSA: The BLSA is supported by the Intramural Research Program, National Institute on Aging, NIH. BLSA is a prospective cohort study with continuous enrollment that began in 1958. Comprehensive data from BLSA are available upon request by a proposal submission through the cohort website (www.blsa.nih.gov).

BIOCARD: The BIOCARD study is designed to identify biomarkers associated with progression from normal cognitive status to cognitive impairment or dementia, with a particular focus on Alzheimer’s Disease. The BIOCARD study is supported by a grant from the National Institute on Aging (NIA): U19-AG03365. The BIOCARD Study consists of 7 Cores and 2 projects with the following members: (1) The Administrative Core (Marilyn Albert, Corinne Pettigrew, Barbara Rodzon); (2) the Clinical Core (Marilyn Albert, Anja Soldan, Rebecca Gottesman, Corinne Pettigrew, Leonie Farrington, Maura Grega, Gay Rudow, Rostislav Brichko, Scott Rudow, Jules Giles, Ned Sacktor); (3) the Imaging Core (Michael Miller, Susumu Mori, Anthony Kolasny,

Hanzhang Lu, Kenichi Oishi, Tilak Ratnanather, Peter vanZijl, Laurent Younes); (4) the Biospecimen Core (Abhay Moghekar, Jacqueline Darrow, Alexandria Lewis, Richard O’Brien); (5) the Informatics Core (Roberta Scherer, Ann Ervin, David Shade, Jennifer Jones, Hamadou Coulibaly, Kathy Moser, Courtney Potter); the (6) Biostatistics Core (Mei-Cheng Wang, Yuxin Zhu, Jiangxia Wang); (7) the Neuropathology Core (Juan Troncoso, David Nauen, Olga Pletnikova, Karen Fisher); (8) Project 1 (Paul Worley, Jeremy Walston, Mei-Fang Xiao), and (9) Project 2 (Mei-Cheng Wang, Yifei Sun, Yanxun Xu.

UK Biobank: This research has been conducted using the UK Biobank resource, application 16315.

CamCAN: Data collection and sharing for this project was provided by the Cambridge Centre for Ageing and Neuroscience (CamCAN). CamCAN funding was provided by the UK Biotechnology and Biological Sciences Research Council (grant number BB/H008217/1), together with support from the UK Medical Research Council and University of Cambridge, UK. Data used in the preparation of this work were obtained from the CamCAN repository (available at http://www.mrc-cbu.cam.ac.uk/datasets/camcan/). The Cambridge Centre for Ageing and Neuroscience (Cam-CAN) is a large-scale collaborative research project at the University of Cambridge.

HBN: The Healthy Brain Network (HBN) is an ongoing initiative focused on building a biobank of data from 10,000 children and adolescents (ages 5-21) in the New York City area (https://www.nature.com/articles/sdata2017181). Data were acquired from the DSI studio website: (https://brain.labsolver.org/hbn.html).

MAP/ROS/MARS: Data contributed from MAP/ROS/MARS was supported by NIA R01AG017917, P30AG10161, P30AG072975, R01AG022018, R01AG056405, UH2NS100599, UH3NS100599, R01AG064233, R01AG15819 and R01AG067482, and the Illinois Department of Public Health (Alzheimer’s Disease Research Fund). Data can be accessed at www.radc.rush.edu. More information about participant demographics and study information can be found here: https://www.rushu.rush.edu/research-rush-university/departmental-research/rush-alzheimers-disease-center/rush-alzheimers-disease-center-research/epidemiologic-research.

The data contributed from the Wisconsin Registry for Alzheimer’s Prevention was supported by NIA AG021155, AG0271761, AG037639, and AG054047.

MORGAN: Data collected from the MORGAN dataset were acquired at Vanderbilt University, with the aim of studying cognitive patterns in epileptic patients both before and after clinical treatment for epilepsy.

CALM: Data collection and sharing for this project was provided by the Centre for Attention, Learning and Memory (CALM). CALM funding was provided by the UK Medical Research Council and University of Cambridge, UK. Data used in the preparation of this work were obtained from CALM resource – https://calm.mrc-cbu.cam.ac.uk/. The study protocol is reported in Holmes et al. (2019).

UTAustin: The dataset that we refer to as UTAustin in this manuscript is a longitudinal neuroimaging dataset on language processing in children ages 5, 7, and 9 years old collected at The University of Texas at Austin. The dataset is openly available from Openneuro here: https://openneuro.org/datasets/ds003604/versions/1.0.7. We use version 1.0.7.

QTAB: The Queensland Twin Adolescent Brain (QTAB) Project was established with the purpose of promoting the conduct of health-related research in adolescence. The QTAB dataset comprises multimodal neuroimaging, as well as cognitive and mental health data collected in adolescent twins over two sessions. The MRI protocol consisted of T1-weighted (MP2RAGE), T2-weighted, FLAIR, high-resolution TSE, SWI, resting-state fMRI, DWI, and ASL scans. The QTAB project resource was produced as a result of i) the goodwill and contribution of 422 twin/triplet participants and their parents, ii) funding from the National Health and Medical Research Council, Australia (APP1078756) and the Queensland Brain Institute, University of Queensland, iii) access to several key resources, including the Centre for Advanced Imaging, the Human Studies Unit, Institute of Molecular Bioscience, and the Queensland Cyber Infrastructure Foundation, at the University of Queensland, local and national twin registries at the QIMR Berghofer Medical Research Institute and Twin Research Australia, as well as the many assessments made available by researchers worldwide, and iv) was established with the purpose of promoting the conduct of health-related research in adolescence. The imaging data and basic demographics are openly accessibly on Openneuro here: https://openneuro.org/datasets/ds004146/versions/1.0.4. We use version 1.0.4.

TempleSocial: The Social Reward and Nonsocial Reward Processing Across the Adult Lifespan: An Interim Multi-echo fMRI and Diffusion Dataset (referred to as TempleSocial in this manuscript) comes from a study that aims to investigate whether older adults have a blunted response to some features of social reward. The dataset is openly accessibly on Openneuro here: https://openneuro.org/datasets/ds005123/versions/1.1.3. In particular, we use version 1.1.3.

Lexical: The Longitudinal Brain Correlates of Multisensory Lexical Processing in Children study (shortened to Lexical in this manuscript) aims to explore developmentally dependent changes in lexical processing for adolescents. The dataset is openly accessibly on Openneuro here: https://openneuro.org/datasets/ds001894/versions/1.4.2. We use version 1.4.2.

Lac5: The UCLA Consortium for Neuropsychiatric Phenomics LA5c Study (UCLA Lac5) is focused on understanding the dimensional structure of memory and cognitive control (response inhibition) functions in both healthy individuals and individuals with neuropsychiatric disorders including schizophrenia, bipolar disorder, and attention deficit/hyperactivity disorder. Neuroimaging data were downloaded from Openneuro here: https://openneuro.org/datasets/ds000030/versions/1.0.0. We use version 1.0.0.

UpennRisk: The dataset we refer to as UPennRisk comes from a study at University of Pennsylvania that investigated whether training executive cognitive function could influence choice behavior and brain responses. Neuroimaging data were downloaded from Openneuro here: https://openneuro.org/datasets/ds002843/versions/1.0.1. We use version 1.0.1.

DLBS: The Dallas Lifespan Brain Study (DLBS) is a longitudinal multi-modal neuroimaging study of the aging mind, which was initiated in 2008 (referred to as Wave 1). Participants returned for two additional waves of data collection with an approximate interval of 4-5 years between waves. The DLBS protocol encompasses various imaging modalities, including structural MRI, diffusion MRI, and functional MRI, as well as comprehensive cognitive and psychosocial assessments. DLBS data can be downloaded from Openneuro here: https://openneuro.org/datasets/ds004856. Specifically, we use version 1.2.0.

MASiVar: The Multisite, Multiscanner, and Multisubject Acquisitions for Studying Variability in Diffusion Weighted Magnetic Resonance Imaging (MASiVar) dataset consists of 319 diffusion scans acquired at 3T from b = 1000 to 3000 s/mm2 across 14 healthy adults, 83 healthy children (5 to 8 years), three sites, and four scanners curated to promote investigation of diffusion MRI variability. In particular, we used only the data coming from healthy children (Cohort IV) for version 2.0.2 of the dataset. Data are available to download from Openneuro here: https://openneuro.org/datasets/ds003416/versions/2.0.2.

SWU: Data coming from the Southwestern University (SWU) dataset, part of the Consortium for Reliability and Reproducibility (CoRR), were downloaded via NITRC-IR from the 1000 Functional Connectomes Project. Specifically, the data come from the Emotion and Creativity One Year Retest Dataset subset, comprised of 235 subjects, all of whom were college students. Each subject underwent two sessions of anatomical, resting state fMRI, and DTI scans, spaced one year apart. In order to access the CoRR datasets through NITRC, users must be logged into NITRC at the time of download and registered with the 1000 Functional Connectomes Project / INDI website. More information about this subset can be found here: https://fcon_1000.projects.nitrc.org/indi/CoRR/html/swu_4.html.

Calgary: The Preschool MRI study in The Developmental Neuroimaging Lab at the University of Calgary (https://www.developmentalneuroimaginglab.ca) uses different magnetic resonance imaging (MRI) techniques to study brain structure and function in early childhood. The study aims to characterize brain development in early childhood, and to offer baseline data that can be used to understand cognitive and behavioural development, as well as to identify deviations from normal development in children with various diseases, disorders, or brain injuries. The MRI techniques used include diffusion tensor imaging (DTI), anatomical imaging, arterial spin labeling (ASL), and resting state functional MRI (rsfMRI). Data can be downloaded from here: https://osf.io/axz5r/.

CUTTING: This work was supported by the National Institute of Child Health and Human Development (NICHD) through awards R01 HD089474 (Cutting), R37 HD095519 (Cutting), R01 HD044073 (Cutting), and R01 HD067254 (Cutting). This work was supported by the Vanderbilt Kennedy Center funded by NICHD through award P50 HD103537. This work utilized REDCap supported by the National Center for Advancing Translational Sciences (NCATS) through award UL1 TR000445 to the Vanderbilt Institute for Clinical and Translational Research. This work was supported by the National Institute of Health’s Office of the Director (1S10 OD021771-01) to the Vanderbilt University Institute of Imaging Science.

AOMIC: The Amsterdam Open MRI Collection (AOMIC) is a collection of three datasets with multimodal (3T) MRI data including structural (T1-weighted), diffusion-weighted, and (resting-state and task-based) functional BOLD MRI data, as well as detailed demographics and psychometric variables from a large set of healthy participants. All raw data is publicly available from the Openneuro data sharing platform: ID1000: https://openneuro.org/datasets/ds003097, PIOP1: https://openneuro.org/datasets/ds002785, PIOP2: https://openneuro.org/datasets/ds002790. We use version 1.2.1 for ID1000 and 2.0.0 for PIOP1 and PIOP2.

BANDA: The Boston Adolescent Neuroimaging of Depression and Anxiety (BANDA) is a study of 215 adolescents ages 14-17, 152 of whom had a current diagnosis of a DSM-5 (APA, 2013) anxious and/or depressive disorder. The BANDA study collected a rich dataset of brain, clinical, and cognitive/neuropsychological measures from these adolescent subjects. The dataset is available to download upon request on the NDA.

ADRC: We thank Knight ADRC for providing neuroimaging data to us. The data contributed through WASHU (Knight ADRC) was supported by grant numbers P30 AG066444, P01 AG03991, and P01 AG026276. For WASHU, Clinical Dementia Ratings (CDRs) are obtained from assessments by experienced clinicians trained in the use of the CDR.

NACC: The NACC database is funded by NIA/NIH Grant U24 AG072122. NACC data are contributed by the NIA-funded ADRCs: P30 AG062429 (PI James Brewer, MD, PhD), P30 AG066468 (PI Oscar Lopez, MD), P30 AG062421 (PI Bradley Hyman, MD, PhD), P30 AG066509 (PI Thomas Grabowski, MD), P30 AG066514 (PI Mary Sano, PhD), P30 AG066530 (PI Helena Chui, MD), P30 AG066507 (PI Marilyn Albert, PhD), P30 AG066444 (PI John Morris, MD), P30 AG066518 (PI Jeffrey Kaye, MD), P30 AG066512 (PI Thomas Wisniewski, MD), P30 AG066462 (PI Scott Small, MD), P30 AG072979 (PI David Wolk, MD), P30 AG072972 (PI Charles DeCarli, MD), P30 AG072976 (PI Andrew Saykin, PsyD), P30 AG072975 (PI David Bennett, MD), P30 AG072978 (PI Ann McKee, MD), P30 AG072977 (PI Robert Vassar, PhD), P30 AG066519 (PI Frank LaFerla, PhD), P30 AG062677 (PI Ronald Petersen, MD, PhD), P30 AG079280 (PI Eric Reiman, MD), P30 AG062422 (PI Gil Rabinovici, MD), P30 AG066511 (PI Allan Levey, MD, PhD), P30 AG072946 (PI Linda Van Eldik, PhD), P30 AG062715 (PI Sanjay Asthana, MD, FRCP), P30 AG072973 (PI Russell Swerdlow, MD), P30 AG066506 (PI Todd Golde, MD, PhD), P30 AG066508 (PI Stephen Strittmatter, MD, PhD), P30 AG066515 (PI Victor Henderson, MD, MS), P30 AG072947 (PI Suzanne Craft, PhD), P30 AG072931 (PI Henry Paulson, MD, PhD), P30 AG066546 (PI Sudha Seshadri, MD), P20 AG068024 (PI Erik Roberson, MD, PhD), P20 AG068053 (PI Justin Miller, PhD), P20 AG068077 (PI Gary Rosenberg, MD), P20 AG068082 (PI Angela Jefferson, PhD), P30 AG072958 (PI Heather Whitson, MD), P30 AG072959 (PI James Leverenz, MD).

SCAN is a multi-institutional project that was funded as a U24 grant (AG067418) by the National Institute on Aging in May 2020. Data collected by SCAN and shared by NACC are contributed by the NIA-funded ADRCs as follows: Arizona Alzheimer’s Center - P30 AG072980 (PI: Eric Reiman, MD); R01 AG069453 (PI: Eric Reiman (contact), MD); P30 AG019610 (PI: Eric Reiman, MD); and the State of Arizona which provided additional funding supporting our center; Boston University - P30 AG013846 (PI Neil Kowall MD); Cleveland ADRC - P30 AG062428 (James Leverenz, MD); Cleveland Clinic, Las Vegas – P20AG068053; Columbia - P50 AG008702 (PI Scott Small MD); Duke/UNC ADRC – P30 AG072958; Emory University - P30AG066511 (PI Levey Allan, MD, PhD); Indiana University - R01 AG19771 (PI Andrew Saykin, PsyD); P30 AG10133 (PI Andrew Saykin, PsyD); P30 AG072976 (PI Andrew Saykin, PsyD); R01 AG061788 (PI Shannon Risacher, PhD); R01 AG053993 (PI Yu-Chien Wu, MD, PhD); U01 AG057195 (PI Liana Apostolova, MD); U19 AG063911 (PI Bradley Boeve, MD); and the Indiana University Department of Radiology and Imaging Sciences; Johns Hopkins - P30 AG066507 (PI Marilyn Albert, Phd.); Mayo Clinic - P50 AG016574 (PI Ronald Petersen MD PhD); Mount Sinai - P30 AG066514 (PI Mary Sano, PhD); R01 AG054110 (PI Trey Hedden, PhD); R01 AG053509 (PI Trey Hedden, PhD); New York University - P30AG066512-01S2 (PI Thomas Wisniewski, MD); R01AG056031 (PI Ricardo Osorio, MD); R01AG056531 (PIs Ricardo Osorio, MD; Girardin Jean-Louis, PhD); Northwestern University - P30 AG013854 (PI Robert Vassar PhD); R01 AG045571 (PI Emily Rogalski, PhD); R56 AG045571, (PI Emily Rogalski, PhD); R01 AG067781, (PI Emily Rogalski, PhD); U19 AG073153, (PI Emily Rogalski, PhD); R01 DC008552, (M.-Marsel Mesulam, MD); R01 AG077444, (PIs M.-Marsel Mesulam, MD, Emily Rogalski, PhD); R01 NS075075 (PI Emily Rogalski, PhD); R01 AG056258 (PI Emily Rogalski, PhD); Oregon Health and Science University - P30 AG008017 (PI Jeffrey Kaye MD); R56 AG074321 (PI Jeffrey Kaye, MD); Rush University - P30 AG010161 (PI David Bennett MD); Stanford – P30AG066515; P50 AG047366 (PI Victor Henderson MD MS); University of Alabama, Birmingham – P20; University of California, Davis - P30 AG10129 (PI Charles DeCarli, MD); P30 AG072972 (PI Charles DeCarli, MD); University of California, Irvine - P50 AG016573 (PI Frank LaFerla PhD); University of California, San Diego - P30AG062429 (PI James Brewer, MD, PhD); University of California, San Francisco - P30 AG062422 (Rabinovici, Gil D., MD); University of Kansas - P30 AG035982 (Russell Swerdlow, MD); University of Kentucky - P30 AG028283-15S1 (PIs Linda Van Eldik, PhD and Brian Gold, PhD); University of Michigan ADRC - P30AG053760 (PI Henry Paulson, MD, PhD) P30AG072931 (PI Henry Paulson, MD, PhD) Cure Alzheimer’s Fund 200775 - (PI Henry Paulson, MD, PhD) U19 NS120384 (PI Charles DeCarli, MD, University of Michigan Site PI Henry Paulson, MD, PhD) R01 AG068338 (MPI Bruno Giordani, PhD, Carol Persad, PhD, Yi Murphey, PhD) S10OD026738-01 (PI Douglas Noll, PhD) R01 AG058724 (PI Benjamin Hampstead, PhD) R35 AG072262 (PI Benjamin Hampstead, PhD) W81XWH2110743 (PI Benjamin Hampstead, PhD) R01 AG073235 (PI Nancy Chiaravalloti, University of Michigan Site PI Benjamin Hampstead, PhD) 1I01RX001534 (PI Benjamin Hampstead, PhD) IRX001381 (PI Benjamin Hampstead, PhD); University of New Mexico - P20 AG068077 (Gary Rosenberg, MD); University of Pennsylvania - State of PA project 2019NF4100087335 (PI David Wolk, MD); Rooney Family Research Fund (PI David Wolk, MD); R01 AG055005 (PI David Wolk, MD); University of Pittsburgh - P50 AG005133 (PI Oscar Lopez MD); University of Southern California - P50 AG005142 (PI Helena Chui MD); University of Washington - P50 AG005136 (PI Thomas Grabowski MD); University of Wisconsin - P50 AG033514 (PI Sanjay Asthana MD FRCP); Vanderbilt University – P20 AG068082; Wake Forest - P30AG072947 (PI Suzanne Craft, PhD); Washington University, St. Louis - P01 AG03991 (PI John Morris MD); P01 AG026276 (PI John Morris MD); P20 MH071616 (PI Dan Marcus); P30 AG066444 (PI John Morris MD); P30 NS098577 (PI Dan Marcus); R01 AG021910 (PI Randy Buckner); R01 AG043434 (PI Catherine Roe); R01 EB009352 (PI Dan Marcus); UL1 TR000448 (PI Brad Evanoff); U24 RR021382 (PI Bruce Rosen); Avid Radiopharmaceuticals / Eli Lilly; Yale - P50 AG047270 (PI Stephen Strittmatter MD PhD); R01AG052560 (MPI: Christopher van Dyck, MD; Richard Carson, PhD); R01AG062276 (PI: Christopher van Dyck, MD); 1Florida - P30AG066506-03 (PI Glenn Smith, PhD); P50 AG047266 (PI Todd Golde MD PhD)

## Supplementary Material

Supplementary figures and tables available online.

## Appendix 1

Group/Consortium Authors

A) HABS-HD MPIs: Sid E O’Bryant, Kristine Yaffe, Arthur Toga, Robert Rissman, & Leigh Johnson; and the HABS-HD Investigators: Meredith Braskie, Kevin King, James R Hall, Melissa Petersen, Raymond Palmer, Robert Barber, Yonggang Shi, Fan Zhang, Rajesh Nandy, Roderick McColl, David Mason, Bradley Christian, Nicole Phillips, Stephanie Large, Joe Lee, Badri Vardarajan, Monica Rivera Mindt, Amrita Cheema, Lisa Barnes, Mark Mapstone, Annie Cohen, Amy Kind, Ozioma Okonkwo, Raul Vintimilla, Zhengyang Zhou, Michael Donohue, Rema Raman, Matthew Borzage, Michelle Mielke, Beau Ances, Ganesh Babulal, Jorge Llibre-Guerra, Carl Hill and Rocky Vig.

B) Data used in preparation of this article were obtained from the Alzheimer’s Disease Neuroimaging Initiative (ADNI) database (adni.loni.usc.edu). As such, the investigators within the ADNI contributed to the design and implementation of ADNI and/or provided data but did not participate in analysis or writing of this report. A complete listing of ADNI investigators can be found at: http://adni.loni.usc.edu/wp-content/uploads/how_to_apply/ADNI_Acknowledgement_List.pdf

C) Data used in preparation of this article were derived from BIOCARD study, supported by grant U19 – AG033655 from the National Institute on Aging. The BIOCARD study team did not participate in the analysis or writing of this report, however, they contributed to the design and implementation of the study. A listing of BIOCARD investigators can be found on the BIOCARD website (on the ‘BIOCARD Data Access Procedures’ page, ‘Acknowledgement Agreement’ document).

## References

Alexander, D. C., Dyrby, T. B., Nilsson, M., & Zhang, H. (2019). Imaging brain microstructure with diffusion MRI: practicality and applications. NMR Biomed, 32(4), e3841. doi:10.1002/nbm.3841

Amemiya, K., Naito, E., & Takemura, H. (2021). Age dependency and lateralization in the three branches of the human superior longitudinal fasciculus. Cortex, 139, 116–133. doi:10.1016/j.cortex.2021.02.027

Amunts, K., Schleicher, A., Burgel, U., Mohlberg, H., Uylings, H. B., & Zilles, K. (1999). Broca’s region revisited: cytoarchitecture and intersubject variability. J Comp Neurol, 412(2), 319–341. doi:10.1002/(sici)1096-9861(19990920)412:2<319::aid-cne10>3.0.co;2-7

Andrulyte, I., De Bezenac, C., Branzi, F., Forkel, S. J., Taylor, P. N., & Keller, S. S. (2024). The Relationship between White Matter Architecture and Language Lateralization in the Healthy Brain. J Neurosci, 44(50). doi:10.1523/JNEUROSCI.0166-24.2024

Angstmann, S., Madsen, K. S., Skimminge, A., Jernigan, T. L., Baaré, W. F., & Siebner, H. R. (2016). Microstructural asymmetry of the corticospinal tracts predicts right-left differences in circle drawing skill in right-handed adolescents. Brain Struct Funct, 221(9), 4475–4489. doi:10.1007/s00429-015-1178-5

Banfi, C., Koschutnig, K., Moll, K., Schulte-Korne, G., Fink, A., & Landerl, K. (2019). White matter alterations and tract lateralization in children with dyslexia and isolated spelling deficits. Hum Brain Mapp, 40(3), 765–776. doi:10.1002/hbm.24410

Beaulieu, C. (2002). The basis of anisotropic water diffusion in the nervous system - a technical review. NMR Biomed, 15(7-8), 435–455. doi:10.1002/nbm.782

Beaulieu, C., Yip, E., Low, P. B., Madler, B., Lebel, C. A., Siegel, L.,…Laule, C. (2020). Myelin Water Imaging Demonstrates Lower Brain Myelination in Children and Adolescents With Poor Reading Ability. Front Hum Neurosci, 14, 568395. doi:10.3389/fnhum.2020.568395

Beck, D., de Lange, A. G., Maximov, II, Richard, G., Andreassen, O. A., Nordvik, J. E., & Westlye, L. T. (2021). White matter microstructure across the adult lifespan: A mixed longitudinal and cross-sectional study using advanced diffusion models and brain-age prediction. Neuroimage, 224, 117441. doi:10.1016/j.neuroimage.2020.117441

Bethlehem, R. A. I., Seidlitz, J., White, S. R., Vogel, J. W., Anderson, K. M., Adamson, C.,…VETSA. (2022). Brain charts for the human lifespan. Nature, 604(7906), 525–533. doi:10.1038/s41586-022-04554-y

Bethlehem, R. A. I., Seidlitz, J., White, S. R., Vogel, J. W., Anderson, K. M., Adamson, C.,…Alexander-Bloch, A. F. (2022). Brain charts for the human lifespan. Nature, 604(7906), 525–533. doi:10.1038/s41586-022-04554-y

Bishop, D. V. (2013). Cerebral asymmetry and language development: cause, correlate, or consequence? Science, 340(6138), 1230531. doi:10.1126/science.1230531

Biswal, B. B., Mennes, M., Zuo, X. N., Gohel, S., Kelly, C., Smith, S. M.,…Milham, M.P. (2010). Toward discovery science of human brain function. Proc Natl Acad Sci U S A, 107(10), 4734–4739. doi:10.1073/pnas.0911855107

Brincat, S. L., & Miller, E. K. (2025). Cognitive independence and interactions between cerebral hemispheres. Neuropsychologia, 212, 109153. doi:10.1016/j.neuropsychologia.2025.109153

Broca, P. (1861). Perte de la parole, ramollissement chronique et destruction partielle du lobe antérieur gauche du cerveau. Bull Soc Anthropol, 2(1), 235–238.

Budisavljevic, S., Castiello, U., & Begliomini, C. (2021). Handedness and White Matter Networks. Neuroscientist, 27(1), 88–103. doi:10.1177/1073858420937657

Cai, L. Y., Yang, Q., Hansen, C. B., Nath, V., Ramadass, K., Johnson, G. W., Landman, B. A. (2021). PreQual: An automated pipeline for integrated preprocessing and quality assurance of diffusion weighted MRI images. Magn Reson Med, 86(1), 456–470. doi:10.1002/mrm.28678

Callaert, D. V., Vercauteren, K., Peeters, R., Tam, F., Graham, S., Swinnen, S. P.,…Wenderoth, N. (2011). Hemispheric asymmetries of motor versus nonmotor processes during (visuo)motor control. Hum Brain Mapp, 32(8), 1311–1329. doi:10.1002/hbm.21110

Carper, R. A., Treiber, J. M., DeJesus, S. Y., & Müller, R. A. (2016). Reduced Hemispheric Asymmetry of White Matter Microstructure in Autism Spectrum Disorder. J Am Acad Child Adolesc Psychiatry, 55(12), 1073–1080. doi:10.1016/j.jaac.2016.09.491

Catani, M., Allin, M. P., Husain, M., Pugliese, L., Mesulam, M. M., Murray, R. M., & Jones, D. K. (2007). Symmetries in human brain language pathways correlate with verbal recall. Proc Natl Acad Sci U S A, 104(43), 17163–17168. doi:10.1073/pnas.0702116104

Catani, M., Craig, M. C., Forkel, S. J., Kanaan, R., Picchioni, M., Toulopoulou, T.,…McGuire, P. (2011). Altered integrity of perisylvian language pathways in schizophrenia: relationship to auditory hallucinations. Biol Psychiatry, 70(12), 1143–1150. doi:10.1016/j.biopsych.2011.06.013

Chen, A., Deng, Y., Zuo, X., & Zhong, S. (2022). Alteration in Asymmetry of White Matter Network of Parkinson’s Disease. Contrast Media Mol Imaging, 2022, 8493729. doi:10.1155/2022/8493729

Claassen, D. O., McDonell, K. E., Donahue, M., Rawal, S., Wylie, S. A., Neimat, J. S.,…Rane, S. (2016). Cortical asymmetry in Parkinson’s disease: early susceptibility of the left hemisphere. Brain Behav, 6(12), e00573. doi:10.1002/brb3.573

Cole, T. J. (2021). Sample size and sample composition for constructing growth reference centiles. Stat Methods Med Res, 30(2), 488–507. doi:10.1177/0962280220958438

Dax, M. (1865). Lesions de la motie gauche de l’encephale coincident avec l’oublie des signes de la pensee. Gaz Hbd Med Chir, 2, 259–262.

Demnitz, N., Madsen, K. S., Johnsen, L. K., Kjaer, M., Boraxbekk, C. J., & Siebner, H. R. (2021). Right-left asymmetry in corticospinal tract microstructure and dexterity are uncoupled in late adulthood. Neuroimage, 240, 118405. doi:10.1016/j.neuroimage.2021.118405

Deoni, S. C., Dean, D. C., 3rd, O’Muircheartaigh, J., Dirks, H., & Jerskey, B. A. (2012). Investigating white matter development in infancy and early childhood using myelin water faction and relaxation time mapping. Neuroimage, 63(3), 1038–1053. doi:10.1016/j.neuroimage.2012.07.037

Dubois, J., Dehaene-Lambertz, G., Kulikova, S., Poupon, C., Huppi, P. S., & Hertz-Pannier, L. (2014). The early development of brain white matter: a review of imaging studies in fetuses, newborns and infants. Neuroscience, 276, 48–71. doi:10.1016/j.neuroscience.2013.12.044

Dubois, J., Grotheer, M., Yang, J. Y., Tournier, J. D., Beaulieu, C., & Lebel, C. (2025). Small brains but big challenges: white matter tractography in early life samples. Brain Struct Funct, 230(4), 58. doi:10.1007/s00429-025-02922-8

Dubois, J., Poupon, C., Thirion, B., Simonnet, H., Kulikova, S., Leroy, F.,…Dehaene-Lambertz, G. (2016). Exploring the Early Organization and Maturation of Linguistic Pathways in the Human Infant Brain. Cereb Cortex, 26(5), 2283–2298. doi:10.1093/cercor/bhv082

Dulyan, L., Bortolami, C., & Forkel, S. J. (2025). Chapter 2 - Asymmetries in the human brain. In C. Papagno&P. Corballis (Eds.), Handbook of Clinical Neurology (Vol. 208, pp. 15–36): Elsevier.

Ford, A., Ammar, Z., Li, L., & Shultz, S. (2023). Lateralization of major white matter tracts during infancy is time-varying and tract-specific. Cereb Cortex, 33(19), 10221–10233. doi:10.1093/cercor/bhad277

Forkel, S. J., Friedrich, P., Thiebaut de Schotten, M., & Howells, H. (2022). White matter variability, cognition, and disorders: a systematic review. Brain Struct Funct, 227(2), 529–544. doi:10.1007/s00429-021-02382-w

Foster, N. N., Barry, J., Korobkova, L., Garcia, L., Gao, L., Becerra, M.,…Dong, H. W. (2021). The mouse cortico-basal ganglia-thalamic network. Nature, 598(7879), 188–194. doi:10.1038/s41586-021-03993-3

Ghasoub, M., Scholten, C., Perdue, M., Long, M., Ostertag, C., Kar, P.,…Lebel, C. (2025). Associations between white matter asymmetry and communication skills in children with prenatal alcohol exposure. Drug Alcohol Depend, 272, 112674. doi:10.1016/j.drugalcdep.2025.112674

Gomez-Gastiasoro, A., Zubiaurre-Elorza, L., Pena, J., Ibarretxe-Bilbao, N., Rilo, O., Schretlen, D. J., & Ojeda, N. (2019). Altered frontal white matter asymmetry and its implications for cognition in schizophrenia: A tractography study. Neuroimage Clin, 22, 101781. doi:10.1016/j.nicl.2019.101781

Good, C. D., Johnsrude, I., Ashburner, J., Henson, R. N., Friston, K. J., & Frackowiak, R. S. (2001). Cerebral asymmetry and the effects of sex and handedness on brain structure: a voxel-based morphometric analysis of 465 normal adult human brains. Neuroimage, 14(3), 685–700. doi:10.1006/nimg.2001.0857

Grech-Sollars, M., Hales, P. W., Miyazaki, K., Raschke, F., Rodriguez, D., Wilson, M.,…Clark, C. A. (2015). Multi-centre reproducibility of diffusion MRI parameters for clinical sequences in the brain. NMR Biomed, 28(4), 468–485. doi:10.1002/nbm.3269

Groen, M. A., Whitehouse, A. J., Badcock, N. A., & Bishop, D. V. (2013). Associations between handedness and cerebral lateralisation for language: a comparison of three measures in children. PLoS One, 8(5), e64876. doi:10.1371/journal.pone.0064876

Grummer-Strawn, L. M., Reinold, C., Krebs, N. F., Centers for Disease, C.,&Prevention. (2010). Use of World Health Organization and CDC growth charts for children aged 0-59 months in the United States. MMWR Recomm Rep, 59(RR-9), 1–15.

Guadalupe, T., Willems, R. M., Zwiers, M. P., Arias Vasquez, A., Hoogman, M., Hagoort, P.,…Francks, C. (2014). Differences in cerebral cortical anatomy of left-and right-handers. Front Psychol, 5, 261. doi:10.3389/fpsyg.2014.00261

Habeck, C., Razlighi, Q., Gazes, Y., Barulli, D., Steffener, J., & Stern, Y. (2017). Cognitive Reserve and Brain Maintenance: Orthogonal Concepts in Theory and Practice. Cereb Cortex, 27(8), 3962–3969. doi:10.1093/cercor/bhw208

Hasan, K. M., Iftikhar, A., Kamali, A., Kramer, L. A., Ashtari, M., Cirino, P. T.,…Ewing-Cobbs, L. (2009). Development and aging of the healthy human brain uncinate fasciculus across the lifespan using diffusion tensor tractography. Brain Res, 1276, 67–76. doi:10.1016/j.brainres.2009.04.025

Hau, J., Sarubbo, S., Houde, J. C., Corsini, F., Girard, G., Deledalle, C.,…Petit, L. (2017). Revisiting the human uncinate fasciculus, its subcomponents and asymmetries with stem-based tractography and microdissection validation. Brain Struct Funct, 222(4), 1645–1662. doi:10.1007/s00429-016-1298-6

Ho, N. F., Li, Z., Ji, F., Wang, M., Kuswanto, C. N., Sum, M. Y.,…Zhou, J. (2017). Hemispheric lateralization abnormalities of the white matter microstructure in patients with schizophrenia and bipolar disorder. J Psychiatry Neurosci, 42(4), 242–251. doi:10.1503/jpn.160090

Honnedevasthana Arun, A., Connelly, A., Smith, R. E., & Calamante, F. (2021). Characterisation of white matter asymmetries in the healthy human brain using diffusion MRI fixel-based analysis. Neuroimage, 225, 117505. doi:10.1016/j.neuroimage.2020.117505

Howells, H., Thiebaut de Schotten, M., Dell’Acqua, F., Beyh, A., Zappala, G., Leslie, A.,…Catani, M. (2018). Frontoparietal Tracts Linked to Lateralized Hand Preference and Manual Specialization. Cereb Cortex, 28(7), 2482–2494. doi:10.1093/cercor/bhy040

Jeurissen, B., Descoteaux, M., Mori, S., & Leemans, A. (2019). Diffusion MRI fiber tractography of the brain. NMR Biomed, 32(4), e3785. doi:10.1002/nbm.3785

Joseph, R. M., Fricker, Z., Fenoglio, A., Lindgren, K. A., Knaus, T. A., & Tager-Flusberg, H. (2014). Structural asymmetries of language-related gray and white matter and their relationship to language function in young children with ASD. Brain Imaging Behav, 8(1), 60–72. doi:10.1007/s11682-013-9245-0

Kim, M. E., Gao, C., Ramadass, K., Kanakaraj, P., Newlin, N. R., Rudravaram, G., OBryant, S. (2024). Scalable quality control on processing of large diffusion-weighted and structural magnetic resonance imaging datasets. arXiv preprint arXiv:2409.17286.

Kim, M. E., Gao, C., Ramadass, K., Newlin, N. R., Kanakaraj, P., Bogdanov, S., Schilling, K. G. (2025). White matter microstructure and macrostructure brain charts across the human lifespan. bioRxiv, 2025.2005.2008.652953. doi:10.1101/2025.05.08.652953

Kim, M. E., Ramadass, K., Gao, C., Kanakaraj, P., Newlin, N. R., Rudravaram, G., Hohman, T. J. (2024). Scalable, reproducible, and cost-effective processing of large-scale medical imaging datasets. arXiv preprint arXiv:2408.14611.

Kong, X. Z., Mathias, S. R., Guadalupe, T., Glahn, D. C., Franke, B., Crivello, F., Group, E. L. W. (2018). Mapping cortical brain asymmetry in 17,141 healthy individuals worldwide via the ENIGMA Consortium. Proc Natl Acad Sci U S A, 115(22), E5154–E5163. doi:10.1073/pnas.1718418115

Kumpulainen, V. (2024). Brain white matter development, associations to maternal perinatal psychological distress and emotional attention at the age of 5 years: fi=Turun yliopisto|en=University of Turku.

Kumpulainen, V., Merisaari, H., Silver, E., Copeland, A., Pulli, E. P., Lewis, J. D.,…Tuulari, J. J. (2023). Sex differences, asymmetry, and age-related white matter development in infants and 5-year-olds as assessed with tract-based spatial statistics. Hum Brain Mapp, 44(7), 2712–2725. doi:10.1002/hbm.26238

Lebel, C., Gee, M., Camicioli, R., Wieler, M., Martin, W., & Beaulieu, C. (2012). Diffusion tensor imaging of white matter tract evolution over the lifespan. Neuroimage, 60(1), 340–352. doi:10.1016/j.neuroimage.2011.11.094

Lebel, C., Treit, S., & Beaulieu, C. (2019). A review of diffusion MRI of typical white matter development from early childhood to young adulthood. NMR Biomed, 32(4), e3778. doi:10.1002/nbm.3778

Manning, L., & Thomas-Anterion, C. (2011). Marc Dax and the discovery of the lateralisation of language in the left cerebral hemisphere. Rev Neurol (Paris), 167(12), 868–872. doi:10.1016/j.neurol.2010.10.017

Marcelle, M., Illapani, V. S. P., Gaillard, W. D., & Newport, E. L. (2025). Assessing the Early Lateralization of White Matter in the Infant Language Network. Hum Brain Mapp, 46(11), e70286. doi:10.1002/hbm.70286

Martel, A. C., & Galvan, A. (2022). Connectivity of the corticostriatal and thalamostriatal systems in normal and parkinsonian states: An update. Neurobiol Dis, 174, 105878. doi:10.1016/j.nbd.2022.105878

Minkova, L., Habich, A., Peter, J., Kaller, C. P., Eickhoff, S. B., & Klöppel, S. (2017). Gray matter asymmetries in aging and neurodegeneration: A review and meta-analysis. Hum Brain Mapp, 38(12), 5890–5904. doi:10.1002/hbm.23772

Miyata, J., Sasamoto, A., Koelkebeck, K., Hirao, K., Ueda, K., Kawada, R.,…Murai, T. (2012). Abnormal asymmetry of white matter integrity in schizophrenia revealed by voxelwise diffusion tensor imaging. Hum Brain Mapp, 33(7), 1741–1749. doi:10.1002/hbm.21326

Mundorf, A., Peterburs, J., & Ocklenburg, S. (2021). Asymmetry in the Central Nervous System: A Clinical Neuroscience Perspective. Front Syst Neurosci, 15, 733898. doi:10.3389/fnsys.2021.733898

Novikov, D. S., Fieremans, E., Jespersen, S. N., & Kiselev, V. G. (2018). Quantifying brain microstructure with diffusion MRI: Theory and parameter estimation. *NMR Biomed*, e3998. doi:10.1002/nbm.3998

O’Muircheartaigh, J., Dean, D. C., 3rd, Dirks, H., Waskiewicz, N., Lehman, K., Jerskey, B. A., & Deoni, S. C. (2013). Interactions between white matter asymmetry and language during neurodevelopment. J Neurosci, 33(41), 16170–16177. doi:10.1523/JNEUROSCI.1463-13.2013

Ocklenburg, S., Anderson, C., Gerding, W. M., Fraenz, C., Schlüter, C., Friedrich, P.,…Genç, E. (2019). Myelin Water Fraction Imaging Reveals Hemispheric Asymmetries in Human White Matter That Are Associated with Genetic Variation in PLP1. Mol Neurobiol, 56(6), 3999–4012. doi:10.1007/s12035-018-1351-y

Ocklenburg, S., Friedrich, P., Güntürkün, O., & Genç, E. (2016). Voxel-wise grey matter asymmetry analysis in left-and right-handers. Neurosci Lett, 633, 210–214. doi:10.1016/j.neulet.2016.09.046

Ocklenburg, S., Hugdahl, K., & Westerhausen, R. (2013). Structural white matter asymmetries in relation to functional asymmetries during speech perception and production. Neuroimage, 83, 1088–1097. doi:10.1016/j.neuroimage.2013.07.076

Ocklenburg, S., Mundorf, A., Gerrits, R., Karlsson, E. M., Papadatou-Pastou, M., & Vingerhoets, G. (2024). Clinical implications of brain asymmetries. Nat Rev Neurol, 20(7), 383–394. doi:10.1038/s41582-024-00974-8

Parekh, S. A., Wren-Jarvis, J., Lazerwitz, M., Rowe, M. A., Powers, R., Bourla, I.,…Mukherjee, P. (2023). Hemispheric lateralization of white matter microstructure in children and its potential role in sensory processing dysfunction. Front Neurosci, 17, 1088052. doi:10.3389/fnins.2023.1088052

Pierpaoli, C., Jezzard, P., Basser, P. J., Barnett, A., & Di Chiro, G. (1996). Diffusion tensor MR imaging of the human brain. Radiology, 201(3), 637–648. doi:10.1148/radiology.201.3.8939209

Powell, H. W., Parker, G. J., Alexander, D. C., Symms, M. R., Boulby, P. A., Wheeler-Kingshott, C. A.,…Duncan, J.S. (2006). Hemispheric asymmetries in language-related pathways: a combined functional MRI and tractography study. Neuroimage, 32(1), 388–399. doi:10.1016/j.neuroimage.2006.03.011

Powell, J. L., Parkes, L., Kemp, G. J., Sluming, V., Barrick, T. R., & García-Fiñana, M. (2012). The effect of sex and handedness on white matter anisotropy: a diffusion tensor magnetic resonance imaging study. Neuroscience, 207, 227–242. doi:10.1016/j.neuroscience.2012.01.016

Propper, R. E., O’Donnell, L. J., Whalen, S., Tie, Y., Norton, I. H., Suarez, R. O.,…Golby, A. J. (2010). A combined fMRI and DTI examination of functional language lateralization and arcuate fasciculus structure: Effects of degree versus direction of hand preference. Brain Cogn, 73(2), 85–92. doi:10.1016/j.bandc.2010.03.004

Rigby, R. A., & Stasinopoulos, D. M. (2005). Generalized Additive Models for Location, Scale and Shape. Journal of the Royal Statistical Society Series C: Applied Statistics, 54(3), 507–554. doi:10.1111/j.1467-9876.2005.00510.x

Rimrodt, S. L., Peterson, D. J., Denckla, M. B., Kaufmann, W. E., & Cutting, L. E. (2010). White matter microstructural differences linked to left perisylvian language network in children with dyslexia. Cortex, 46(6), 739–749. doi:10.1016/j.cortex.2009.07.008

Roe, J. M., Vidal-Pineiro, D., Amlien, I. K., Pan, M., Sneve, M. H., Thiebaut de Schotten, M.,…Westerhausen, R. (2023). Tracing the development and lifespan change of population-level structural asymmetry in the cerebral cortex. Elife, 12. doi:10.7554/eLife.84685

Roll, M. (2024). Heschl’s gyrus and the temporal pole: The cortical lateralization of language. Neuroimage, 303, 120930. doi:10.1016/j.neuroimage.2024.120930

Saltoun, K., Yeo, B. T. T., Paul, L., Diedrichsen, J., & Bzdok, D. (2025). Longitudinal changes in brain asymmetry track lifestyle and disease. Nat Commun, 16(1), 5611. doi:10.1038/s41467-025-60451-8

Santoni, G., Calderon-Larranaga, A., Vetrano, D. L., Welmer, A. K., Orsini, N., & Fratiglioni, L. (2020). Geriatric Health Charts for Individual Assessment and Prediction of Care Needs: A Population-Based Prospective Study. J Gerontol A Biol Sci Med Sci, 75(1), 131–138. doi:10.1093/gerona/gly272

Savic, I., & Lindström, P. (2008). PET and MRI show differences in cerebral asymmetry and functional connectivity between homo-and heterosexual subjects. Proc Natl Acad Sci U S A, 105(27), 9403–9408. doi:10.1073/pnas.0801566105

Schilling, K. G., Chad, J. A., Chamberland, M., Nozais, V., Rheault, F., Archer, D.,…Landman, B. A. (2023). White matter tract microstructure, macrostructure, and associated cortical gray matter morphology across the lifespan. Imaging Neuroscience, 1, 1–24. doi:10.1162/imag_a_00050

Schilling, K. G., Rheault, F., Petit, L., Hansen, C. B., Nath, V., Yeh, F. C.,…Descoteaux, M. (2021). Tractography dissection variability: What happens when 42 groups dissect 14 white matter bundles on the same dataset? Neuroimage, 243, 118502. doi:10.1016/j.neuroimage.2021.118502

Schilling, K. G., Tax, C. M. W., Rheault, F., Hansen, C. B., Yang, Q., Yeh, F.-C.,…Landman, B. A. (2021). Fiber tractography bundle segmentation depends on scanner effects, acquisition, diffusion sensitization, and bundle segmentation workflow. bioRxiv, 2021.2003.2017.435872. doi:10.1101/2021.03.17.435872

Schilling, K. G., Tax, C. M. W., Rheault, F., Landman, B. A., Anderson, A. W., Descoteaux, M., & Petit, L. (2021). Prevalence of white matter pathways coming into a single white matter voxel orientation: The bottleneck issue in tractography. Hum Brain Mapp. doi:10.1002/hbm.25697

Schmithorst, V. J., Holland, S. K., & Dardzinski, B. J. (2008). Developmental differences in white matter architecture between boys and girls. Hum Brain Mapp, 29(6), 696–710. doi:10.1002/hbm.20431

scilus/scilpy. The sherbrooke connectivity imaging lab (scil) python dmri processing tool-box. In.

Shepherd, G. M. G., & Yamawaki, N. (2021). Untangling the cortico-thalamo-cortical loop: cellular pieces of a knotty circuit puzzle. Nat Rev Neurosci, 22(7), 389–406. doi:10.1038/s41583-021-00459-3

Shu, M., Yu, C., Shi, Q., Li, Y., Niu, K., Zhang, S., & Wang, X. (2021). Alterations in white matter integrity and asymmetry in patients with benign childhood epilepsy with centrotemporal spikes and childhood absence epilepsy: An automated fiber quantification tractography study. Epilepsy Behav, 123, 108235. doi:10.1016/j.yebeh.2021.108235

Shu, N., Liu, Y., Duan, Y., & Li, K. (2015). Hemispheric Asymmetry of Human Brain Anatomical Network Revealed by Diffusion Tensor Tractography. Biomed Res Int, 2015, 908917. doi:10.1155/2015/908917

Sporns, O., Tononi, G., & Kotter, R. (2005). The human connectome: A structural description of the human brain. PLoS Comput Biol, 1(4), e42. doi:10.1371/journal.pcbi.0010042

Takao, H., Hayashi, N., & Ohtomo, K. (2013). White matter microstructure asymmetry: effects of volume asymmetry on fractional anisotropy asymmetry. Neuroscience, 231, 1–12. doi:10.1016/j.neuroscience.2012.11.038

Thiebaut de Schotten, M., Dell’Acqua, F., Forkel, S. J., Simmons, A., Vergani, F., Murphy, D. G., & Catani, M. (2011). A lateralized brain network for visuospatial attention. Nat Neurosci, 14(10), 1245–1246. doi:10.1038/nn.2905

Thiebaut de Schotten, M., Ffytche, D. H., Bizzi, A., Dell’Acqua, F., Allin, M., Walshe, M.,…Catani, M. (2011). Atlasing location, asymmetry and inter-subject variability of white matter tracts in the human brain with MR diffusion tractography. Neuroimage, 54(1), 49–59. doi:10.1016/j.neuroimage.2010.07.055

Toga, A. W., & Thompson, P. M. (2003). Mapping brain asymmetry. Nat Rev Neurosci, 4(1), 37–48. doi:10.1038/nrn1009

Villalón-Reina, J. E., Zhu, A. H., Benavidez, S., Moreau, C. A., Feng, Y., Chattopadhyay, T.,…Initiative, t. A. s. D. N. (2024). Lifespan Normative Modeling of Brain Microstructure. bioRxiv, 2024.2012.2015.628527. doi:10.1101/2024.12.15.628527

Vingerhoets, G., Verhelst, H., Gerrits, R., Badcock, N., Bishop, D. V. M., Carey, D.,…consortium, L. (2023). Laterality indices consensus initiative (LICI): A Delphi expert survey report on recommendations to record, assess, and report asymmetry in human behavioural and brain research. Laterality, 28(2-3), 122–191. doi:10.1080/1357650X.2023.2199963

Wasserthal, J., Neher, P., & Maier-Hein, K. H. (2018). TractSeg - Fast and accurate white matter tract segmentation. Neuroimage, 183, 239–253. doi:10.1016/j.neuroimage.2018.07.070

Wheeler-Kingshott, C. A., & Cercignani, M. (2009). About “axial” and “radial” diffusivities. Magn Reson Med, 61(5), 1255–1260. doi:10.1002/mrm.21965

Wiberg, A., Ng, M., Al Omran, Y., Alfaro-Almagro, F., McCarthy, P., Marchini, J.,…Furniss, D. (2019). Handedness, language areas and neuropsychiatric diseases: insights from brain imaging and genetics. Brain, 142(10), 2938–2947. doi:10.1093/brain/awz257

Wolff, S. B. E., Ko, R., & Ölveczky, B. P. (2022). Distinct roles for motor cortical and thalamic inputs to striatum during motor skill learning and execution. Sci Adv, 8(8), eabk0231. doi:10.1126/sciadv.abk0231

Wu, Y., Sun, D., & Wang, Y. (2016). Subcomponents and Connectivity of the Inferior Fronto-Occipital Fasciculus Revealed by Diffusion Spectrum Imaging Fiber Tracking. Front Neuroanat, 10, 88. doi:10.3389/fnana.2016.00088

Yeh, F. C. (2020). Shape analysis of the human association pathways. Neuroimage, 223, 117329. doi:10.1016/j.neuroimage.2020.117329

Zhu, A. H., Nir, T. M., Javid, S., Villalon-Reina, J. E., Rodrigue, A. L., Strike, L. T.,…Initiative, A. s. D. N. (2024). Lifespan reference curves for harmonizing multi-site regional brain white matter metrics from diffusion MRI. bioRxiv, 2024.2002.2022.581646. doi:10.1101/2024.02.22.581646

Zhu, Y., Li, S., Da, X., Lai, H., Tan, C., Liu, X.,…Chen, L. (2023). Study of the relationship between onset lateralization and hemispheric white matter asymmetry in Parkinson’s disease. J Neurol, 270(10), 5004–5016. doi:10.1007/s00415-023-11849-1

