## Supplementary Information for "Lifespan Trajectories of Asymmetry in White Matter Tracts"

**Handedness-stratified normative modeling**

Handedness was available for a subset of 12 cohorts (N = 14,220). To assess whether tract-level asymmetry differed by handedness, we repeated the normative modeling framework within this subset and estimated separate LI trajectories for right- and left-handed individuals for exemplar pathways and features (Supplementary Figure 1). Across tract–feature combinations, handedness-related differences in the modeled median LI were consistently small (Cohen’s d typically < 0.08, frequently < 0.02), indicating minimal handedness dependence of structural asymmetry in this dataset. Because handedness was not available across the full consortium and left-handed individuals were relatively sparse and unevenly distributed across the lifespan, these curves are presented as supplementary/exploratory analyses rather than as primary reference charts.

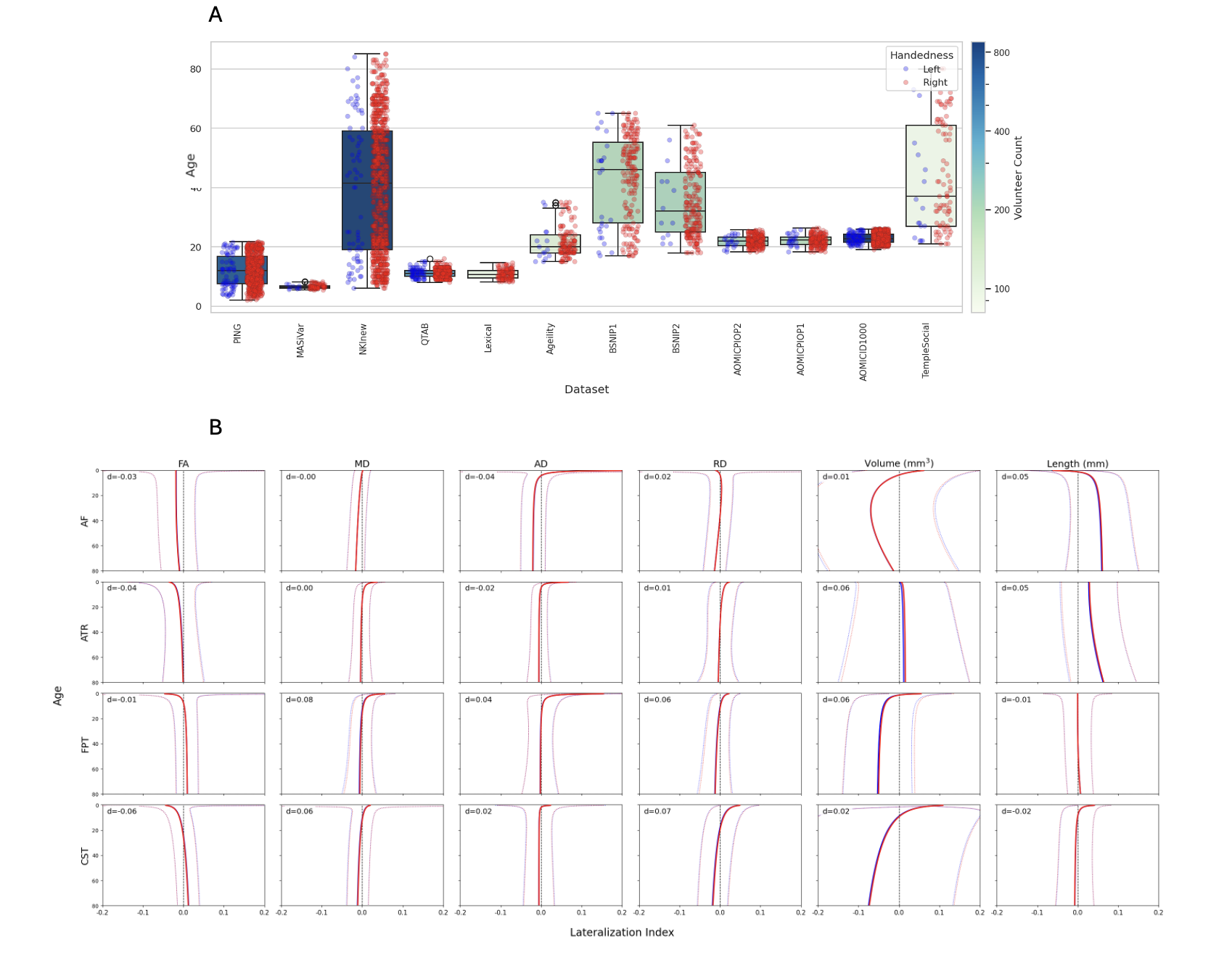

**Supplementary Figure 1: Handedness effects on lifespan white matter asymmetry are small.** Handedness-stratified normative trajectories of the laterality index (LI) are shown for right-handed (blue) and left-handed (red) individuals in the subset of cohorts with handedness data (12 cohorts; N = 15,548; ambidextrous individuals excluded from the handedness-stratified curves). Rows correspond to example tracts (AF, ATR, FPT, CST) and columns to features (FA, MD, AD, RD, tract volume, tract length). Solid lines denote the modeled population median (50th centile) LI as a function of age; dotted lines indicate the normative spread (2.5th and 97.5th centiles). The vertical dashed line marks LI = 0 (symmetry); positive values indicate rightward asymmetry and negative values indicate leftward asymmetry. Panel annotations report Cohen’s d, computed as the mean separation between handedness-specific median LI trajectories across age (standardized by pooled variability), and demonstrate consistently small handedness-related differences across tract–feature combinations.

**Complete lifespan LI trajectories**

To complement the exemplar plots shown in the main text, we provide the full set of sex-stratified lifespan trajectories for lateralization indices across all tracts and measures, enabling visual comparison of asymmetry magnitude and age-dependence across the entire white matter connectome.

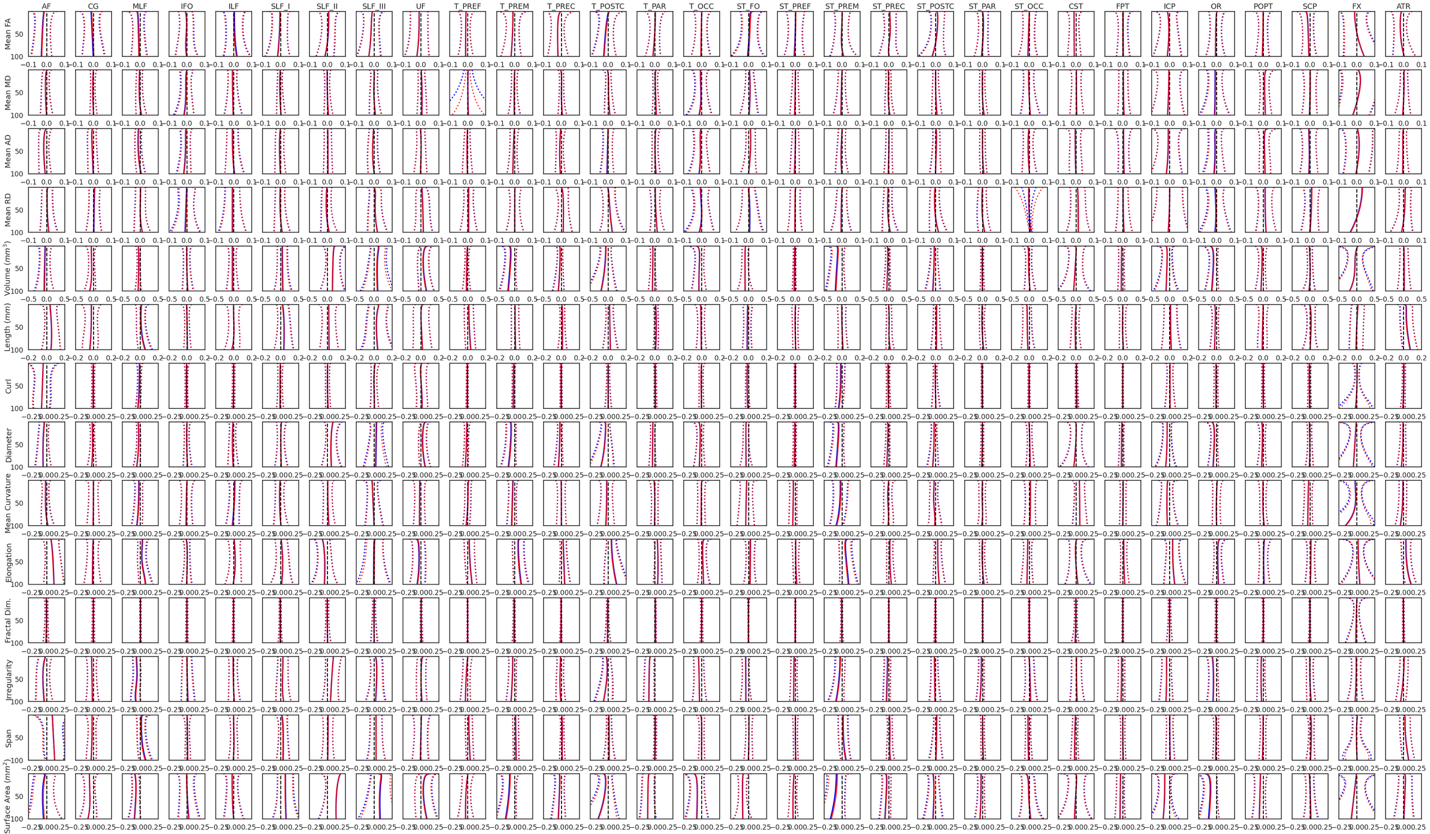

**Supplementary Figure 2: Age-related trajectories of lateralization indices across white matter tracts and diffusion/shape measures**. Each subplot displays the median (solid lines) and 95% confidence interval (dotted lines) of lateralization index (LI) across age for males (blue) and females (red). LI was modeled using the GAMLSS framework with a normal distribution family. Rows represent different microstructural and geometric measures (e.g., FA, MD, volume, curvature), while columns correspond to specific white matter tracts (e.g., AF, SLF, CST). Negative values indicate leftward asymmetry; positive values indicate rightward asymmetry. Vertical dashed lines at x = 0 highlight the zero-asymmetry point. Curves are aligned across tracts to enable direct comparison of asymmetry magnitude and lifespan trajectory by sex and measure. We observe asymmetry patterns change across tracts, measure, and age.

**When asymmetry changes most rapidly**

Because visually similar LI trajectories can reflect different developmental timing, we also show the first derivative of the median LI curves for representative tracts and measures to highlight periods of accelerated change and relative stability across the lifespan.

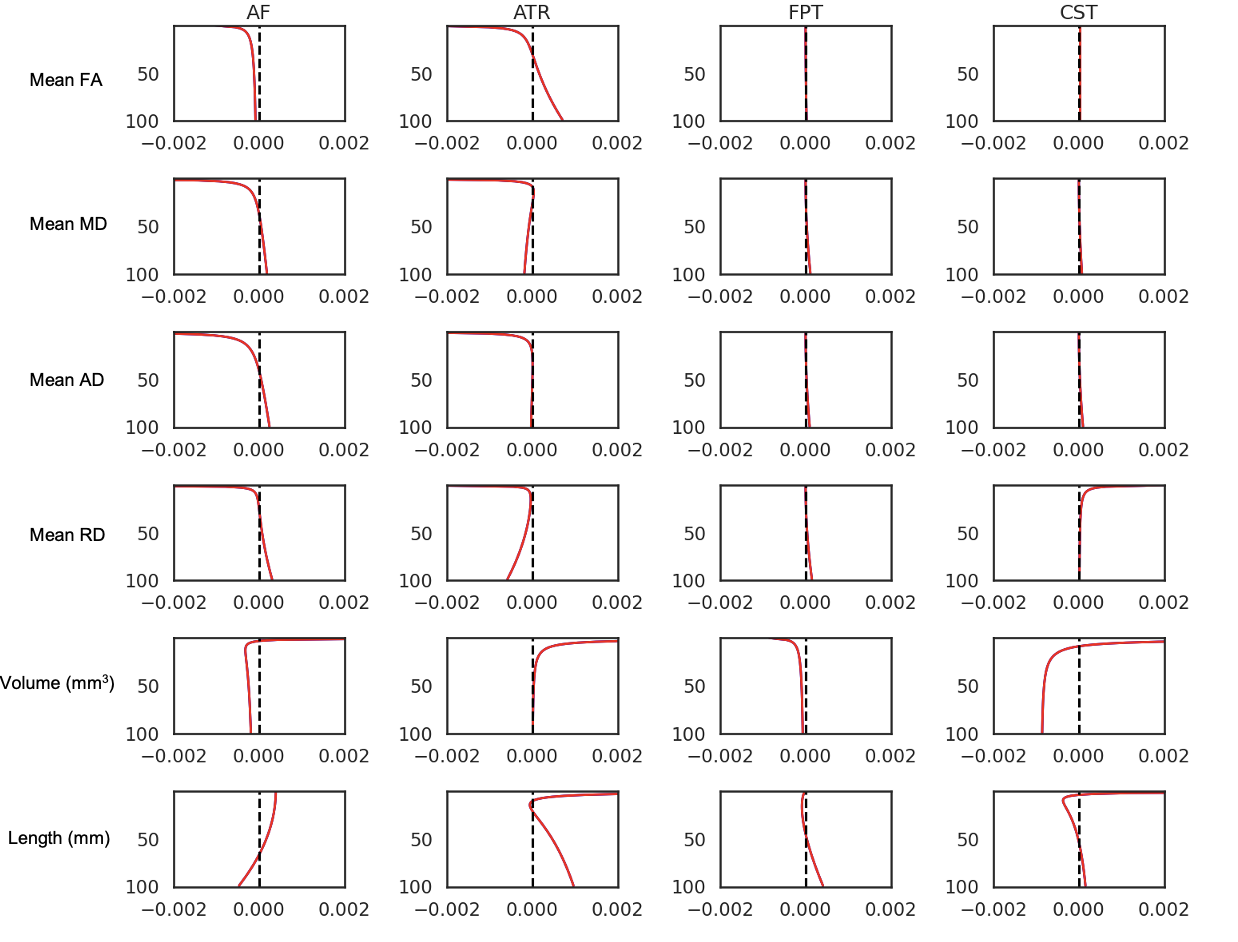

**Supplementary Figure 3. Age-related derivatives of the median LI across white matter tracts and measures.** Each subplot shows the first derivative of the 50th percentile (median) LI trajectory across age (y-axis, from 1 to 100 years) for both males (blue) and females (red) (visually hard to see as they overlap each other). Columns correspond to four major white matter tracts (AF, ATR, FPT, CST), and rows correspond to six asymmetry measures: fractional anisotropy (FA), mean diffusivity (MD), axial diffusivity (AD), radial diffusivity (RD), volume, and average streamline length. The vertical dashed line at x = 0 indicates zero change in asymmetry. Derivative curves show strong early developmental changes, especially in diffusion measures, with greater variability in structural metrics and some divergence between sexes.

**Cohort composition after QC**

To document the contributing data underlying the normative models, we summarize scan and participant counts for each cohort before and after quality control, along with the neurotypical subset retained for analysis.

**Supplementary Table 1: Overview of 50 Neuroimaging Datasets*.*** Including Subject, Session, and Scan Totals

| **Dataset** | **Scans** | **Participants** | **Participants After Quality Control** | **Neurotypical** |
| --- | --- | --- | --- | --- |
| ABCD | 20951 | 10607 | 10311 | 9000 |
| ADNI | 3451 | 1292 | 1229 | 584 |
| AOMIC PIOP1 | 211 | 211 | 204 | 204 |
| AOMIC PIOP2 | 226 | 226 | 224 | 224 |
| AOMIC ID1000 | 925 | 925 | 858 | 858 |
| Ageility | 131 | 131 | 131 | 131 |
| BANDA | 207 | 207 | 207 | 63 |
| BIOCARD | 974 | 210 | 186 | 153 |
| BLSA | 5423 | 1035 | 995 | 942 |
| BSNIP1 | 1400 | 594 | 554 | 197 |
| BSNIP2 | 1432 | 730 | 718 | 233 |
| Calgary | 249 | 88 | 88 | 88 |
| CALM | 314 | 314 | 313 | 189 |
| CAMCAN | 641 | 641 | 613 | 613 |
| VUMC-ASD | 173 | 164 | 132 | 63 |
| CUTTING | 1661 | 638 | 632 | 597 |
| dHCP | 753 | 668 | 285 | 285 |
| DLBS | 955 | 464 | 444 | 444 |
| EBDT | 1389 | 468 | 455 | 455 |
| HABSHD | 3767 | 2762 | 2558 | 1885 |
| HBCD | 546 | 500 | 285 | 285 |
| HBN | 2760 | 2754 | 2519 | 1049 |
| HCPA | 719 | 719 | 718 | 718 |
| HCPBaby | 411 | 211 | 197 | 197 |
| HCPD | 621 | 621 | 621 | 621 |
| HCP | 1109 | 1065 | 1064 | 1064 |
| ABC-Babies | 180 | 142 | 85 | 85 |
| IBIS | 494 | 272 | 217 | 167 |
| ICBM | 264 | 192 | 185 | 185 |
| Lexical | 113 | 113 | 112 | 112 |
| MAP | 1579 | 589 | 393 | 273 |
| MARS | 347 | 184 | 172 | 143 |
| MASiVar | 118 | 83 | 83 | 83 |
| MORGAN | 414 | 256 | 254 | 110 |
| NACC | 1768 | 1353 | 677 | 485 |
| NKI | 2224 | 1289 | 1205 | 840 |
| PING | 762 | 762 | 695 | 695 |
| QTAB | 717 | 415 | 414 | 414 |
| ROS | 127 | 77 | 46 | 35 |
| SCAN | 303 | 288 | 260 | 188 |
| SWU | 460 | 234 | 231 | 231 |
| TempleSocial | 108 | 108 | 103 | 103 |
| UCLA | 262 | 262 | 248 | 116 |
| UKBB | 10425 | 10425 | 8509 | 8509 |
| UPennRisk | 251 | 152 | 151 | 151 |
| UTAustin579 | 376 | 239 | 237 | 237 |
| VMAP_2.0 | 342 | 327 | 266 | 266 |
| VMAP | 1239 | 328 | 310 | 160 |
| TN Alz Project | 194 | 168 | 159 | 58 |
| WRAP | 570 | 343 | 338 | 332 |
| ***Total*** | ***75036*** | ***46846*** | ***41891*** | ***35120*** |

**Diffusion acquisition heterogeneity across cohorts**

Because dMRI measures can vary with acquisition protocol, we report voxel size and diffusion weighting/sampling (b-values and directions) for each cohort, together with scanning location, to contextualize cross-site variability in the aggregated dataset.

**Supplemental Table 2.** Acquisition information for dMRI data from each dataset, along with the respective scanning/facility location.

| **Dataset** | **Voxel Size** | **Acquisition Parameters** | **Facility Location** |
| --- | --- | --- | --- |
| ABCD[1] | 1.7 x 1.7 x 1.7 | 7 b0; 6 b500; 15 b1000; 15 b2000; 60 b3000 | Varied (USA) |
| ADNI[2] | 2.0 x 2.0 x 2.0 | (Varied) 7 b0; 48 b1000 | Varied (USA) |
| Ageility[3] | 2.0 x 2.0 x 2.0 | 1 b0; 64 b1000; 64 b3000 | Newcastle, Australia |
| AOMIC ID1000[4] | 2.0 x 2.0 x 2.0 | 1 b0; 32 b1000 [sequence acquired x3] | Amsterdam, Netherlands |
| AOMIC PIOP1[4] | 2.0 x 2.0 x 2.0 | 1 b0; 32 b1000 | Amsterdam, Netherlands |
| AOMIC PIOP2[4] | 2.0 x 2.0 x 2.0 | 1 b0; 32 b1000 | Amsterdam, Netherlands |
| BANDA[5] | 1.5 x 1.5 x 1.5 | 28 b0; 190 b1500; 190 b3000 | Boston, Massachusetts (USA) |
| BIOCARD[6] | 0.828 x 0.828 x 2.2 | 1 b0; 32 b700 | Baltimore, Maryland (USA) |
| BLSA[7] | 0.8125 x 0.8125 x 2.2^$^ | 1 b0; 32 b700 | Baltimore, Maryland (USA) |
| BSNIP1[8] | 1.72 x 1.72 x 3.0 | 1 b0; 32 b1000 | Varied (USA) |
| BSNIP2[9] | 2.0 x 2.0 x 2.0 | 7 b0; 64 b900 | Varied (USA) |
| Calgary[10] | 0.78 x 0.78 x 2.2 | 5 b0; 30 b800 | Calgary, Canada |
| CALM[11] | 2.0 x 2.0 x 2.0 | 5 b0; 64 b1000 | Cambridge, England |
| CAMCAN[12] | 2.0 x 2.0 x 2.0 | 3 b0; 30 b1000; 30 b2000 | Cambridge, England |
| VUMC-ASD | 2.5 x 2.5 x 2.5 | 1 b0; 92 b1600 3 b0; 6 b500; 32 b1000; 64 b2000 | Nashville, Tennessee (USA) |
| CUTTING | 2.5 x 2.5 x 2.5 | 1 b0; 60 b2000 | Nashville, Tennessee (USA) |
| dHCP[13] | 1.5 x 1.5 x 1.5 | 20 b0; 64 b400; 88 b1000; 128 b2600 | United Kingdom |
| DLBS[14] | 1.75 x 1.75 x 3.0 | 1 b0; 30 b1000 | Dallas, Texas (USA) |
| EBDT[15] | 2.0 x 2.0 x 2.0 | 7 b0; 42 b1000 | Chapel Hill, North Carolina (USA) |
| HABSHD[16] | 1.72 x 1.72 x 2.5 | 4 b0; 64 b1000 [sequence acquired x3] | Fort Worth, Texas (USA) |
| HBCD[17] | 1.7 x 1.7 x 1.7 | 22 b0; 12 b500; 24 b1000; 36 b2000; 58 b3000 | Varied (USA) |
| HBN[18] | 1.8 x 1.8 x1.8 | 1 b0; 64 b1000; 64 b2000 | Varied (USA) |
| HCP[19] | 1.25 x 1.25 x 1.25 | 18 b0; 90 b1000; 90 b2000; 90 b3000 | St. Louis, Missouri; Minneapolis, Michigan (USA) |
| HCPA[20] | 1.5 x 1.5 x 1.5 | 28 b0; 186 b1500; 184 b3000 | St. Louis, Missouri; Minneapolis, Michigan (USA) |
| HCPBaby[21] | 1.5 x 1.5 x 1.5 | 7 b0; 18 b500; 24 b1000; 108 b1500; 48 b2000; 68 b2500; 170 b3000 | Chapel Hill, North Carolina; Minneapolis, Michigan (USA) |
| HCPD[22] | 1.5 x 1.5 x 1.5 | 28 b0; 186 b1500; 184 b3000 | St. Louis, Missouri; Minneapolis, Michigan (USA) |
| ABC-Babies | 1.78 x 1.78 x2.2 | 1 b0; 32 b700; 64 b2000 | Nashville, Tennessee (USA) |
| IBIS[23] | 2.0 x 2.0 x 2.0 | 1 b0; multi-shell (b=100-1000 in steps of 100), 2-3 volumes/shell | Varied (USA) |
| ICBM[24] | 1.25 x 1.25 x 2.5 | 5 b0; 30 b1000 | Los Angeles, California (USA) |
| Lexical[25] | 2.0 x 2.0 x 2.0 | 1 b0; 64 b1000 | Chicago, Illinois (USA) |
| MASiVar[26] | 2.14 x 2.14 x 2.2 | 34 b0; 86 b1000; 112 b2000 | Boston, Massachusetts (USA) |
| MORGAN | 2.5 x 2.5 x 2.5 | 1 b0; 92 b1600 | Nashville, Tennessee (USA) |
| NACC[27] | 2.0 x 2.0 x 2.0 | (Varied) 1 b0; 64 b1000 | Varied (USA) |
| NKI[28] | 2.0 x 2.0 x 2.0 | 9 b0; 128 b1500 | Rockland County, New York (USA) |
| PING[29] | 2.5 x 2.5 x 2.5 | 1 b0; 32 b1000 | Varied (USA) |
| QTAB[30] | 2.0 x 2.0 x 2.0 | 3 b0; 5 b1000; 15 b3000 | Queensland, Australia |
| ROS[31] | 2.0 x 2.0 x 2.0 | 6 b0; 40 b1000 | Chicago, Illinois (USA) |
| MAP[32] | 2.0 x 2.0 x 2.0 | 6 b0; 40 b1000 | Chicago, Illinois (USA) |
| MARS[33] | 2.0 x 2.0 x 2.0 | 1 b0; 40 b1000 | Chicago, Illinois (USA) |
| SCAN[34] | 2.0 x 2.0 x 2.0 | 13 b0; 6 b500; 48 b1000; 60 b2000 | Varied (USA) |
| SWU[35] | 2.0 x 2.0 x 2.0 | 3 b0; 30 b1000 | Chongqing, China |
| TempleSocial[36,37] | 2.0 x 2.0 x 2.0 | 3 b0; 6 b300; 21 b1000; 24 b2000; 12 b3200; 19 b3300; 61 b5000 | Philadelphia, Pennsylvania (USA) |
| UCLA_LA5c[38] | 1.98 x 1.98 x2.0 | 1 b0; 64 b1000 | Los Angeles, California (USA) |
| UKBB[39] | 2.02 x 2.02 x 2 | 8 b0; 50 b1000; 50 b2000 | United Kingdom |
| UPennRisk[40,41] | 1.875 x 1.875 x 2.0 | 1 b0; 30 b1000 | Philadelphia, Pennsylvania (USA) |
| UTAustin579[42] | 2.0 x 2.0 x 4.0 | 6 b0; 64 b800 | Austin, Texas (USA) |
| VMAP_2.0[43] | 2.33 x 2.33 x 2.5 | 1 b0; 38 b1000; 56 b2000 | Nashville, Tennessee (USA) |
| VMAP[43] | 2.0 x 2.0 x 2.0 | 1 b0; 32 b1000 | Nashville, Tennessee (USA) |
| TN Alzheimer’s Project | 2.33 x 2.33 x 2.5 | 1 b0; 38 b1000; 56 b2000 | Nashville, Tennessee (USA) |
| WRAP[44] | 0.9375 x 0.9375 x 2.5 | 8 b0; 40 b1300 | Madison, Wisconsin ((USA) |

**Prior literature context**

To situate the present work within the existing evidence base, we summarize prior studies of white matter asymmetry across different age ranges, including pathways/features examined, sample size, and primary findings.

**Supplementary Table 2. Review of studies investigating white matter asymmetry at various points in the lifespan*.*** The study method, micro- and macro-structural features investigated, number of participants, age range, and key findings are summarized. ^8,17,18,20-23,26,45-47,50,55,57,58,63-83^

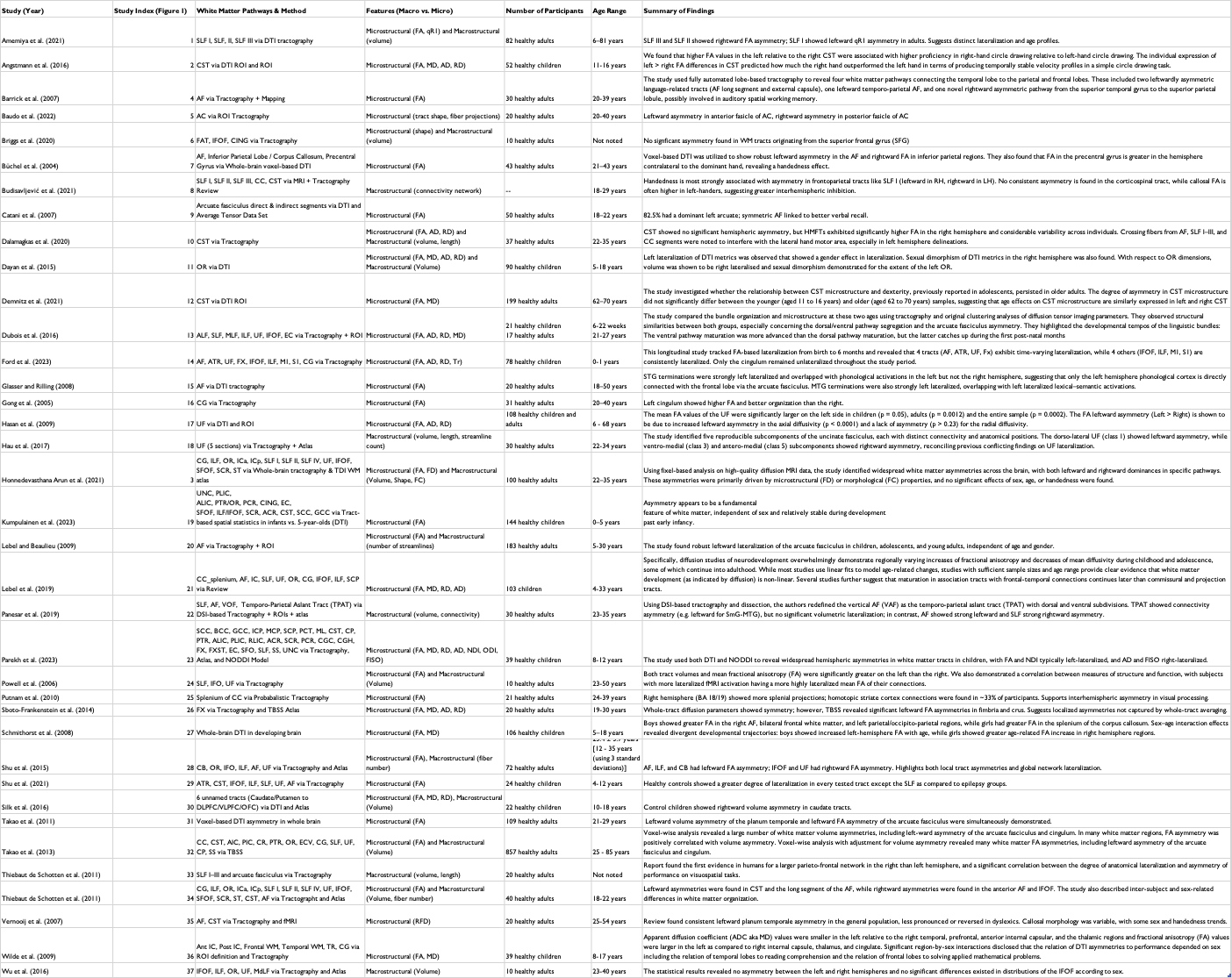
